# Mathematical Modeling Identifies the Role of Adaptive Immunity as a Key Controller of Respiratory Syncytial Virus (RSV) Titer in Cotton Rats

**DOI:** 10.1101/451302

**Authors:** Darren Wethington, Olivia Harder, Karthik Uppulury, William C. L. Stewart, Phylip Chen, Tiffany King, Susan D. Reynolds, Mark E. Peeples, Stefan Niewiesk, Jayajit Das

**Author notes:** Equal contribution (Darren Wethington, Olivia Harder).

## Abstract

Respiratory syncytial virus (RSV) is a common virus that can have varying effects ranging from mild cold-like symptoms to mortality depending on the age and immune status of the individual. We combined mathematical modeling using ordinary differential equations (ODEs) with measurement of RSV infection kinetics in primary well differentiated human airway epithelial (HAE) cultures *in vitro* and in immunocompetent and immunosuppressed cotton rats to glean mechanistic details that underlie RSV infection kinetics in the lung. Quantitative analysis of viral titer kinetics in our mathematical model showed that the elimination of infected cells by the adaptive immune response generates unique RSV titer kinetic features including a faster time scale of viral titer clearance than viral production, and a monotonic decrease in the peak RSV titer with decreasing inoculum dose. Parameter estimation in the ODE model using a non-linear mixed effects approach revealed a very low rate (average single cell lifetime > 10 days) of cell lysis by RSV before the adaptive immune response is initiated. Our model predicted negligible changes in the RSV titer kinetics on earlier days (< 5 d.p.i) but a slower decay in RSV titer at later days (>5 d.p.i) in immunosuppressed cotton rats compared to that in non-suppressed cotton rats *in silico.* These predictions were in excellent agreement with the experimental results. Our combined approach quantified the importance of the adaptive immune response in suppressing RSV infection in cotton rats, which could be useful in testing RSV vaccine candidates.

**Importance:** A major difficulty in developing vaccines against RSV infection is our rudimentary understanding of the mechanisms that underlie RSV infection. We addressed this challenge by developing a mechanistic computational model with predictive powers for describing RSV infection kinetics in cotton rats. The model was constructed synergistically with *in vitro* and *in vivo* measurements. The combined framework determined an important role for CD8+ T cells responses in reducing RSV titers in cotton rats. The framework can be used to design future experiments to elucidate mechanisms underlying RSV infection and test outcomes for potential vaccine candidates. In addition, estimation of the model parameters provides quantitative values for parameters of biological and clinical interest such as the replication rate of RSV, the death rate of infected cells, and the average number of new infections initiated by a single infected cell.

## Introduction

Almost every individual is infected with respiratory syncytial virus (RSV) by age two. Though RSV infection usually does not cause any concern in healthy adults, it can cause serious morbidity and even death in children^1^, immune-compromised individuals^2^, and the elderly^2^. RSV infection is a major cause of hospital admissions and death worldwide in children under five years of age^3^. Despite the public health relevance of RSV infection there is no vaccine or anti-viral therapy available and treatment is supportive^4^.

The RSV virion is surrounded by a lipid membrane with three glycoproteins. The attachment (G) glycoprotein is important for infection but the fusion (F) glycoprotein is essential. RSV infects the ciliated epithelial cells lining of the airway in the respiratory tract by first attaching to the host cell plasma membrane via the G protein and then fusing its membrane with the membrane of the cell, a process mediated by the F protein^5,6^. Once the RSV nucleocapsid enters the cytoplasm, its polymerase produces mRNA and replicates its genome, using the host cell machinery to produce proteins and to assemble virions that are released into the local environment^5^.

RSV owes its middle name to the RSV F protein that also causes fusion of infected cells with neighboring cells, resulting in large cells (syncytia) containing multiple nuclei. Syncytia are more common in RSV-infected immortalized cells in tissue culture^7^ than *in vivo*^8^. Both the innate and adaptive immune responses of the host limit RSV infection but the effectiveness of the adaptive response depends on the age of the individual^9^. The innate response is mediated by interferons generated by the epithelial cells, macrophages, neutrophils, natural killer (NK) cells, and maternal antibodies in neonates^10^. The adaptive immune response, usually induced several days (≥ 4 days) after the infection, is mediated by cytolytic CD8+ T cells, helper CD4+ T cells, and B cells^9^.

The specific roles of the components of the innate and adaptive immune response in controlling RSV infection are not well understood. Challenges of investigating these aspects in the high-risk group, namely neonates, immune-compromised adults, and the elderly contribute to this problem. This presents a major roadblock in developing vaccines and anti-viral therapies.

Animal models such as cotton rats infected with lab strains of RSV have been successfully employed to analyze roles of immune response in regulating RSV infection^11,12^. To this end, we combined mathematical models^13-16^ based on population dynamics of viruses and the host immune response with viral titer measurements in cotton rats to determine the roles of the adaptive and innate immune responses in controlling the viral titer. Population dynamics models have been successfully employed for describing kinetics of human immunodeficiency virus (HIV)^13,17,18^, hepatitis C virus (HCV)^19^, and influenza A virus (IAV)^14,15,20^ infections within the host and some of these models have been used to develop vaccination strategies against these infections. Few population dynamic models have been developed to describe RSV infection kinetics *in vitro*^21^ and *in vivo*^22^. These studies^21,22^ employ viral kinetic models developed for IAV infection for describing RSV kinetics. However, RSV and IAV infection within the host differ in several key aspects, such as IAV infection self-limiting by cytopathic death of target cells^8,14^, whereas RSV induces few obvious cytopathic effects *in vivo*^8,23^. In addition, the adaptive immune response, an important regulator of RSV infection, is not considered in these models. The model developed here shows the importance of the adaptive immune response in generating key features of RSV infection in cotton rats which cannot be captured by simple extensions of IAV models. We performed a detailed parameter estimation using a non-linear mixed effects modeling approach^16,24^.

Furthermore, we tested model predictions affirmatively with experiments carried out in CD8+ T cell depleted and cyclophosphamide treated cotton rats. Our results demonstrate the role of the adaptive immune response; in particular, lysis of the RSV-infected epithelial cells by CD8+ T cells.

## Results

### Evaluation of distinct features of RSV titer kinetics

We used RSV titer measurements in cotton rats inoculated with RSV as reported by Prince et al.^12^ to evaluate relevant time scales and to characterize the shape of the viral titer kinetics. We analyzed the kinetic trajectories followed by the geometric mean of the RSV titers measured in three animals^12^. The following time scales were calculated (Fig. 1A): (1) Peak time (τ_peak_) or the time where the viral titer reached its maximum (or peak) value (V_peak_) post infection, (2) production time (τ_prod_) or the time duration where the viral titer increased *f* times to reach the peak value, (3) decay time (τ_decay_) or the time duration where the viral titer decreased *f* times from the peak value. A dimensionless variable τ_ratio_ = τ_decay_/τ_prod_ characterizes the shape of the RSV viral titer kinetics, i.e., τ_ratio_ <1 indicates a slower time scale for the production of RSV in the early stages (t≤ τ_peak_) of the infection compared to the time scale of reduction of the viral load by the host immune response after the peak titer is reached, whereas, τ_ratio_ >1 marks the opposite scenario. We found that the RSV titers measured in the lungs of cotton rats at different ages (3 days (neonate), 14 days, 28 days or 6 to 8 weeks (adult)) produced τ_ratio_ ≤1 (feature#1) (Fig. 1B). This behavior appears to be unique to RSV infection as influenza A virus (IAV) titer kinetics^25^ in the lung of cotton rats displays τ_ratio_ >1 (see Fig. S1 for an analysis of uncertainties in τ_ratio_ estimation). The IAV titer in the lungs of the cotton rats, unlike inbred laboratory mouse and humans, becomes undetectable in ∼1 day due to the presence of antiviral Mx protein^26^ in the cytoplasm in cotton rats. This specific feature of RSV titer kinetics appears to be organ dependent as the RSV titer kinetics in the nose of the same cohort of cotton rats displayed a τ_ratio_ ≥ 1 at several ages of the animal cohorts.

**Fig. 1.**
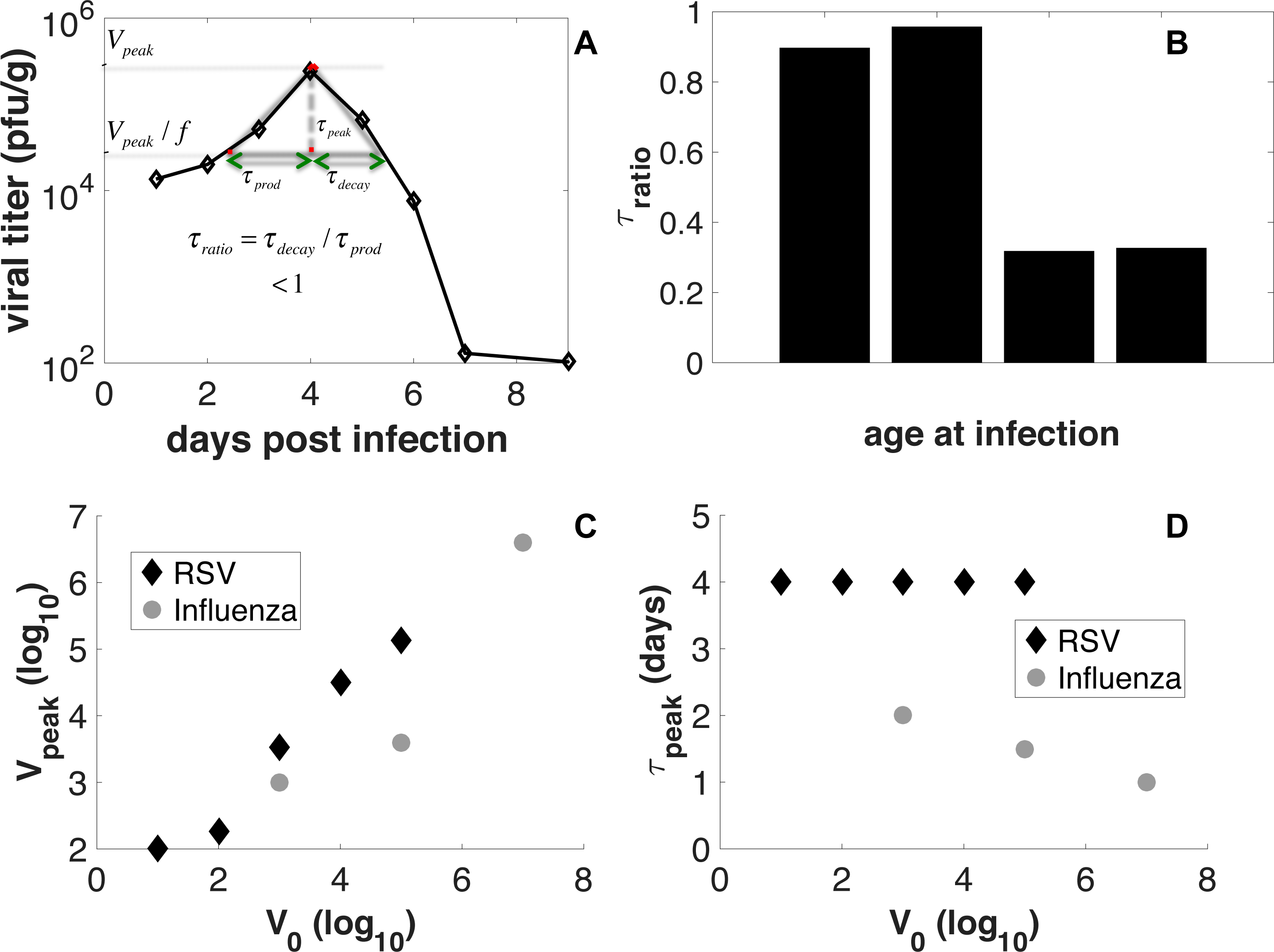
Shape features of RSV titer kinetics. (A) Schematic diagram defining τ_prod_, τ_decay_, τ_peak_, and τ_ratio_. (B) Calculation of τ_ratio_ for the RSV titer kinetics in the lung of cotton rats reported in Prince et al.^12^ (C) Variation of Vpeak with V_0_ for RSV (black diamonds) and influenza A infection (grey circles) in cotton rats. (D) Variation of Vpeak with V_0_ for RSV (black diamonds) and influenza A infection (grey circles) in cotton rats.

Next we analyzed the variation of τ_peak_ and V_peak_ with increasing inoculation dose (Vinocul) in cotton rats. Several experiments show τ_peak_ for RSV infection occurs at roughly 4 days post infection irrespective of the of the Vinocul dose over a 100 fold range (feature #2) (Fig. 1C). Additionally, experiments by Prince and colleagues suggested V_peak_ increased monotonically with increasing V_inocul_ (feature #3) (Fig. 1D)^27^. For influenza A infection in cotton rats, τ_peak_ decreased but V_peak_ increased with increasing Vinocul for the viral loads measured in the lungs (Figs. 1C-D)^25^. Thus, some of the above features (feature #1 and feature #2) appear to be unique to RSV infection in the lungs of cotton rats.

We set out to develop a minimal model that is able to capture the above features of RSV kinetics. As a caveat, we note there are several features of the RSV titer kinetics we do not attempt to model here, e.g., the RSV titer in the lung of the cotton rats show the presence of plateaus or multiple peaks in the 28 day old or adult cotton rats^12^. The presence of these characteristics is more common in the RSV kinetics in the trachea (see Fig. 2 in Ref. ^12^) and in the nose (Fig. 3 in Ref. ^12^). These features could arise due to the interplay between RSV titer and Type I interferons (IFNs) that inhibit RSV replication (see Discussion). Interferons are induced within hours of RSV infection as a component of host innate immunity. Therefore, capturing these features in a mathematical model will require a more complicated model with many unknown parameters. Since measurements of viral titers at a few time points are available, it is likely that multiple complex models with parameters lying in wide ranges of possible values will be able to explain the data. This will make drawing mechanistic conclusions from the analysis difficult. Thus here we pursue modeling few broad features as described above for the RSV titer kinetics with a minimal model.

**Fig. 2.**
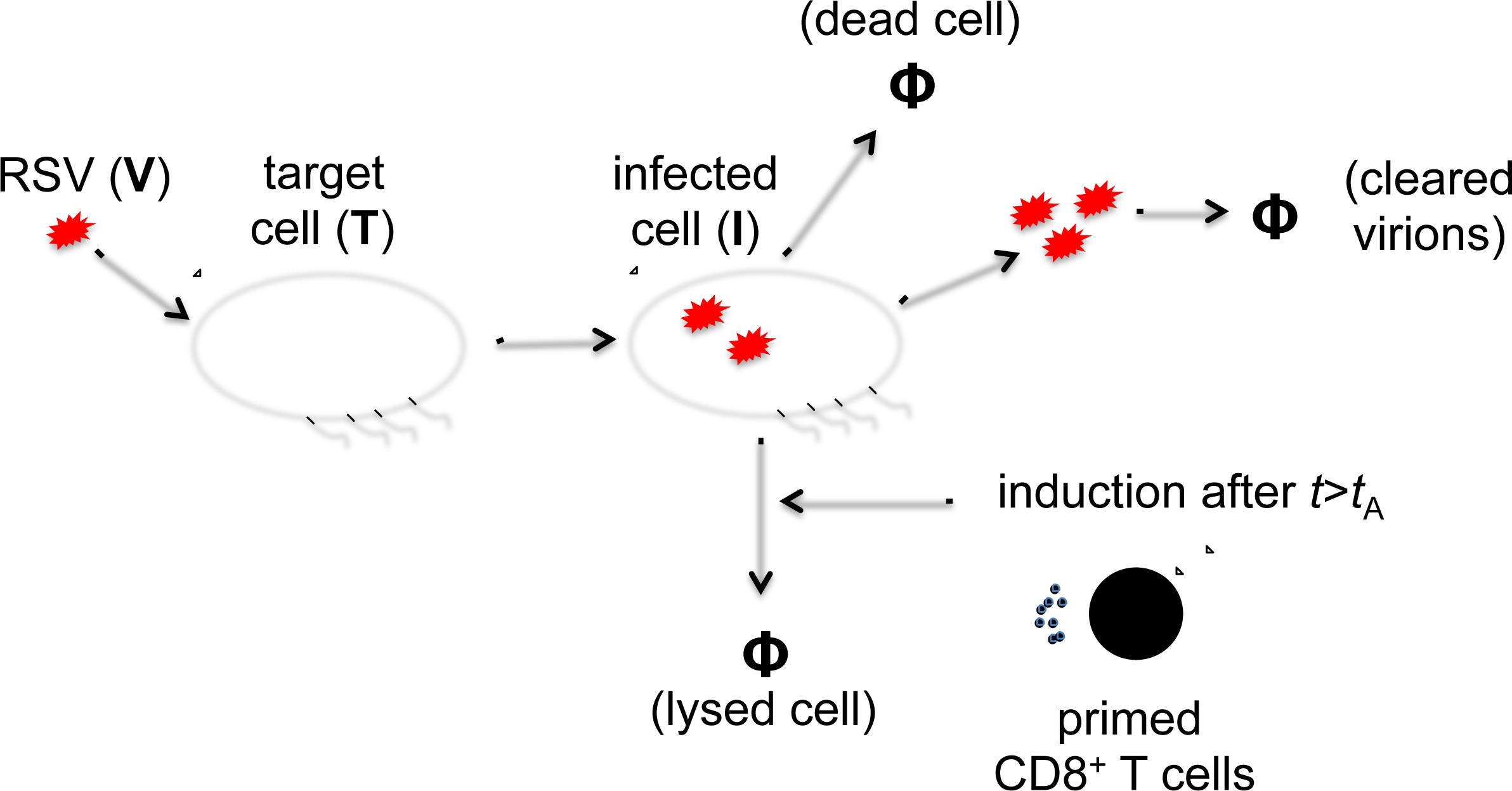
Schematic diagram for the processes described in the RSV model. Ciliated target cells (T) are infected by RSV virions (V) by a rate (*βTV*) producing infected cells (I). The variables (e.g., T, I, V) used in the ODEs for describing the biological entities are denoted in regular fonts and their abundances are indicated in italics (e.g., *T*, *I*, *V*). The infected cells produce new RSV virions at a rate (*pI*), and undergo apoptosis at a rate *δI* due to RSV infection. Free virions are cleared at a rate *cV* from the local environment due to mucociliary activity or lose infection capability. After a time lag t_A_ post infection primed CD8+ T cells lyse RSV infected cells with a rate *δ^(I)^ IA*. The CD8+ T cell response (*A*) is modeled by a step function which changes from 0 to 1 at *t=t_A_*.

**Fig. 3.**
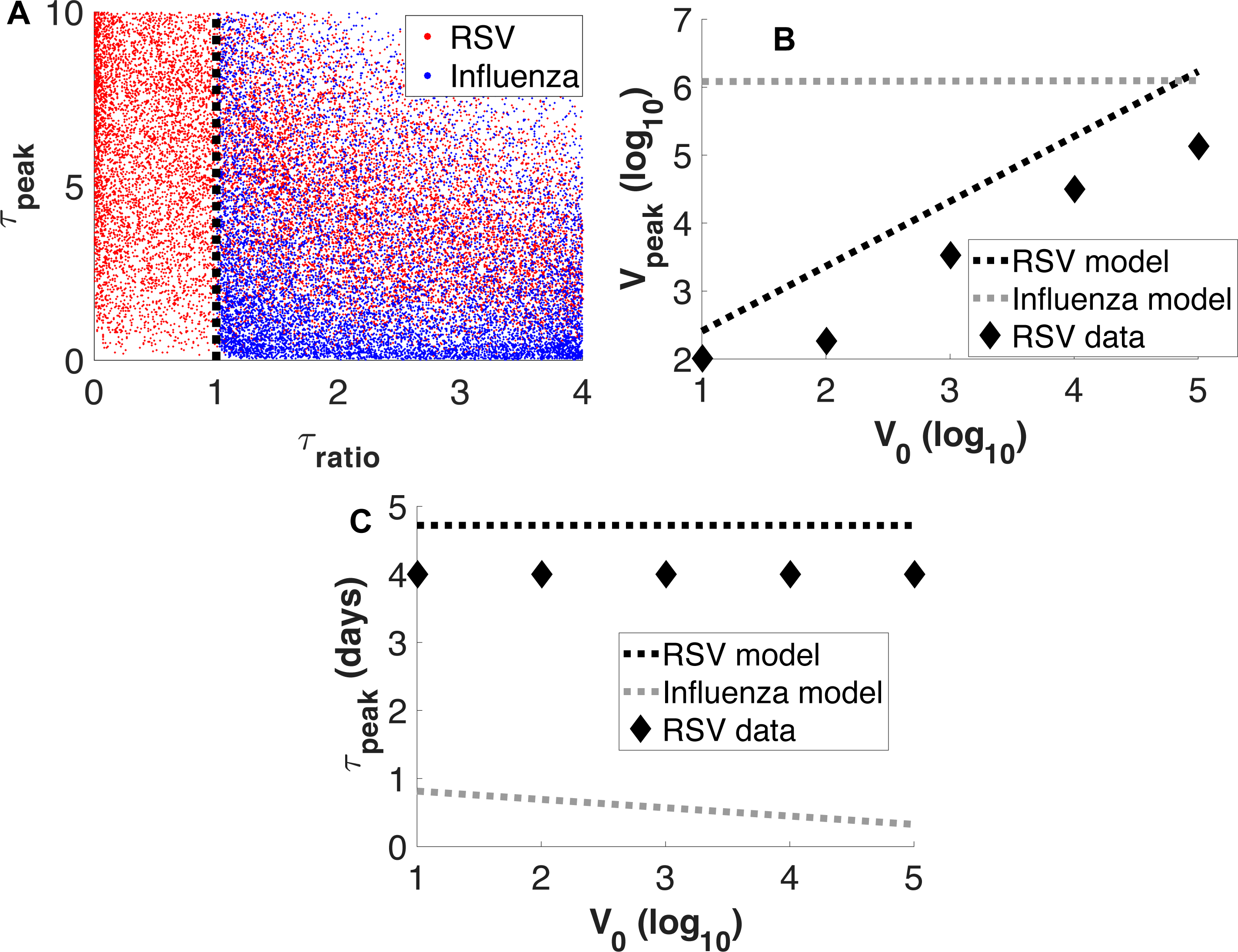
The adaptive immune response (δ^(I)^≠0) in the minimal model is essential for generating the unique features of RSV titer kinetics. (A) Values of τ_ratio_ and τ_peak_ obtained for the viral titer kinetics in the ODE model with (δ^(I)^ ≠0; RSV model) and without (δ^(I)^ = 0; IAV model) the adaptive immune response. 26,854 number of cases for the RSV model and 34,861 number of cases for the IAV model are shown. (B) Variation of V_peak_ with V_0_ in the ODE model with (δ^(I)^ ≠0; RSV model) and without (δ^(I)^ =0; IAV model) the adaptive immune response. (C) Variation of τ_peak_ with V_0_ in the ODE model with (δ^(I)^ ≠0; RSV model) and without (δ^(I)^ = 0; IAV model) the adaptive immune response.

### Mathematical modeling of RSV titer kinetics

We developed a minimal model (Fig. 2A) to describe kinetics of RSV titer in cotton rats in terms of the number of ciliated target cells (*T*), RSV infected cells (*I*), and free RSV virions (*V*). In the model, ciliated target cells become infected by the RSV virions^8,28^ to produce infected cells at rate *βTV*. The infected cells then produce and release RSV virions at a rate *pI*into the local environment. The infected cells are destroyed by RSV at a rate *δI* and RSV virions are cleared at a rate *cV* from the local environment (Fig. 2). The adaptive immune response (e.g., CD8^+^ T cell response) *A* is modeled as a binary variable which changes from zero to unity at *t=t_A_*. The adaptive immune response eliminates the infected cells with a rate *δ*^(*I*)^*I A*. These processes are given by the following set of coupled ordinary differential equations (ODEs).

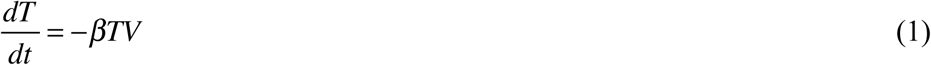

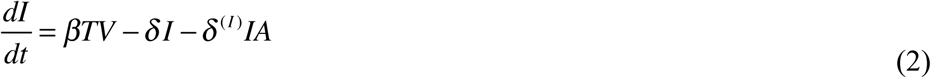

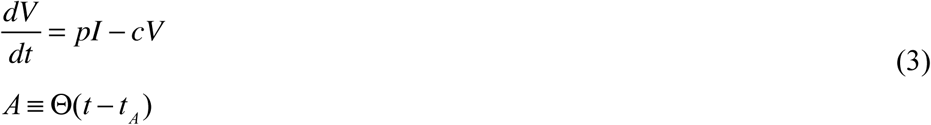

The above ODEs minimally describe attachment and entry of RSV virions in the ciliated target cells followed by production and release of the RSV virions by the infected cells. Experiments using infection of human epithelial cells in tissue cultures suggest RSV to be mildly cytopathic implying smaller values for the rate δ^8^.

The adaptive immunity against RSV infection is complex and is mediated by T cell and B cell responses where these immune cells are activated by RSV antigens presented by antigen presenting cells. Activated CD8^+^ T cells clear the infection by directly eliminating infected epithelial cells. In addition, CD4^+^ T cells and regulatory T cells participate in limiting the infection by influencing B cell responses and controlling the cytokine milieu^9^. In cotton rats CD8^+^ and CD4^+^ T cell responses are initiated around 4 days post infection. Activated B cells secrete antibodies to neutralize the virus, however, these neutralizing antibodies come into action in cotton rats at later days (≥6 dpi in experiments by Prince et al^12^) when the RSV titers are already receding. In our experiments the neutralizing antibodies were detected around day 12 when the RSV titers reached very low or undetectable values.

In addition, we found that depletion of CD4+ T cells in our experiments does not change viral clearance indicating that during the acute phase the CD4+ T cell response does not play a substantial role in clearance (Table. S1). Therefore, we assumed only the CD8+ T cell response in the minimal model. The nature of CD8+ T cell response regulated by CD4+ T cells, B cells, and cytokines can be complex^9^, however, we modeled the response minimally by a binary variable *A* that is turned on at a time t_A_ post infection. This is done in the spirit of finding the simplest model that can be used to describe the viral titer kinetics by approximating the kinetics of many other parts of the immune and host response associated with the infection. In addition, maternal antibodies^9^ were not present in the cotton rats in our experiments. The other innate immune responses mediated by interferons, and NK cells were not accounted for in the model for simplicity. We also note that the same ODEs in Eqs. (1)-(3) with δ^(I)^=0 have been used to model influenza A kinetics in humans^14^, however, the term describing the adaptive immune response (=δ^(I)^IA) plays a key role in capturing the specific features of RSV titer kinetics (see the next section for details).

### Elimination of the infected cells by the CD8^+^ T cell response in the model is necessary for generating key features of RSV infection kinetics

We report our analysis regarding the ability of the model in Eqs. (1)-(3) to describe the features of the RSV titer kinetics qualitatively here and describe the parameter estimation in the next section. The parameters (β, δ, δ^(I)^, t_A_, p, c, and V_0_) in the ODE model (Eqs. (1)-(3)) were varied 10,000 fold across base values (Table S2). We found that the presence of the CD8^+^ T cell induced elimination of the infected cells is important for generating these features. In the absence of this response (or δ^(I)^=0) the RSV titer kinetics were limited to the region τ_ratio_ ≥1(Fig. 3A). Note, when δ^(I)^=0, the resulting ODE model is the same as that used by Baccam et al.^14^ to describe influenza A infection in humans. In contrast, for δ^(I)^≠0, τ_ratio_ spanned a wider region (0< τ_ratio_ ≤1 and τ_ratio_ >1) which included the measured values of τ_ratio_ and the peak time for the RSV titer kinetics in cotton rats (Fig. 3A). Furthermore, the increase of the peak RSV titer V_peak_ and the weak dependency of the peak time τ_peak_ with increasing inoculum dose (V_0_) was captured by the RSV kinetics for δ^(I)^ ≠0 (Figs. 3B-C). However, when δ^(I)^ was set to zero in the model keeping the other parameters the same, the model failed to describe these features qualitatively. In this case, V_peak_ did not change appreciably and τ_peak_ decreased as V_0_ increased (Figs. 3B-C).

### Modeling kinetics of RSV infection of human airway epithelial (HAE) cultures grown *in vitro*

The modeling of RSV infection kinetics in HAE cultures *in vitro* was carried out to achieve three main goals: (1) Evaluate how well the ODE model without the adaptive immune response (δ^(I)^=0) was able to capture the *in vitro*kinetics. (2) Estimate parameters in the model. Since the *in vitro* measurements performed here are able to measure kinetics of both numbers of infected cells (I) and RSV titer (V) at many time points and control the inoculation dose (V_0_) and the number of target cells (T_0_), these measurements provide well defined data to constrain parameter estimates in the model. (3) Use the estimated parameters as an initial guess for the model parameters for fitting the ODE model to the RSV titer measurements in cotton rats. The parameter estimation for the *in vivo* model was carried out using a non-linear mixed effects modeling approach which involves optimization of cost functions in large dimensions, therefore, the parameter estimations are often sensitive to the initial guesses and the ranges of the parameter values^16,24^. We will assume that the common model parameters estimated in the *in vitro* experiments for HAE cultures provide biologically relevant values and therefore use these values as initial guesses for estimating parameters for RSV infection in the cotton rat.

The HAE cells were grown in a tissue culture and were infected with a human strain of recombinant RSV expressing green fluorescent protein (rgRSV)^8^. The tissue culture was washed with 100 µL of medium every 24 hours and the RSV titer in the 100µL of washed medium is measured (see Materials and Methods for details). We set δ^(I)^=0 for the RSV infection model describing the *in vitro* infection since there is no adaptive immune response in this case. Eq. (3) was modified to account for the periodic removal of RSV titer where the RSV titer V(t) was reduced by a fixed fraction (λ) of V(t) at t=t _r_≡{24h, 48h, 72h, 96h, and so on}. The viral kinetics for t_r_ + Δt≤t_r_+24h (Δ → 0) was evaluated by numerically solving Eqs. (1)-(3) for δ^(I)^=0 with initial conditions at t=t_r_ +Δ as T(t_r_+Δ)=T(t_r_), I(t_r_+Δ)=I(t_r_), and V(t_r_+Δ)=V(t_r_) - λ V(t_r_) (see Fig. S2 for further details). λV(t_r_) and I(t_r_) were fitted with their experimental counter parts over a span of 14 days post infection (Fig. 4). The fits to the kinetics of RSV titer and the infected cells were excellent (Fig. 4). The model parameters were estimated using the method of nonlinear least squares where the sum of the squares of the difference between the values obtained for the model simulation and *in vitro* measurements corresponding to logarithms of the viral titer and the number of infected cells was minimized. The calculations were performed using the *lsqcurvefit* subroutine in MATLAB using the Levenberg-Marquardt algorithm. The parameter values, the confidence intervals, and the reproductive number R_0_ =β*pT*_0_/(*c*δ) are shown in Table I for λ=0.3, where the cost function was the minimum (Fig. S3). R_0_ provides an estimate of the average number of new infections initiated by a single infected cell^29^.

**Fig. 4.**
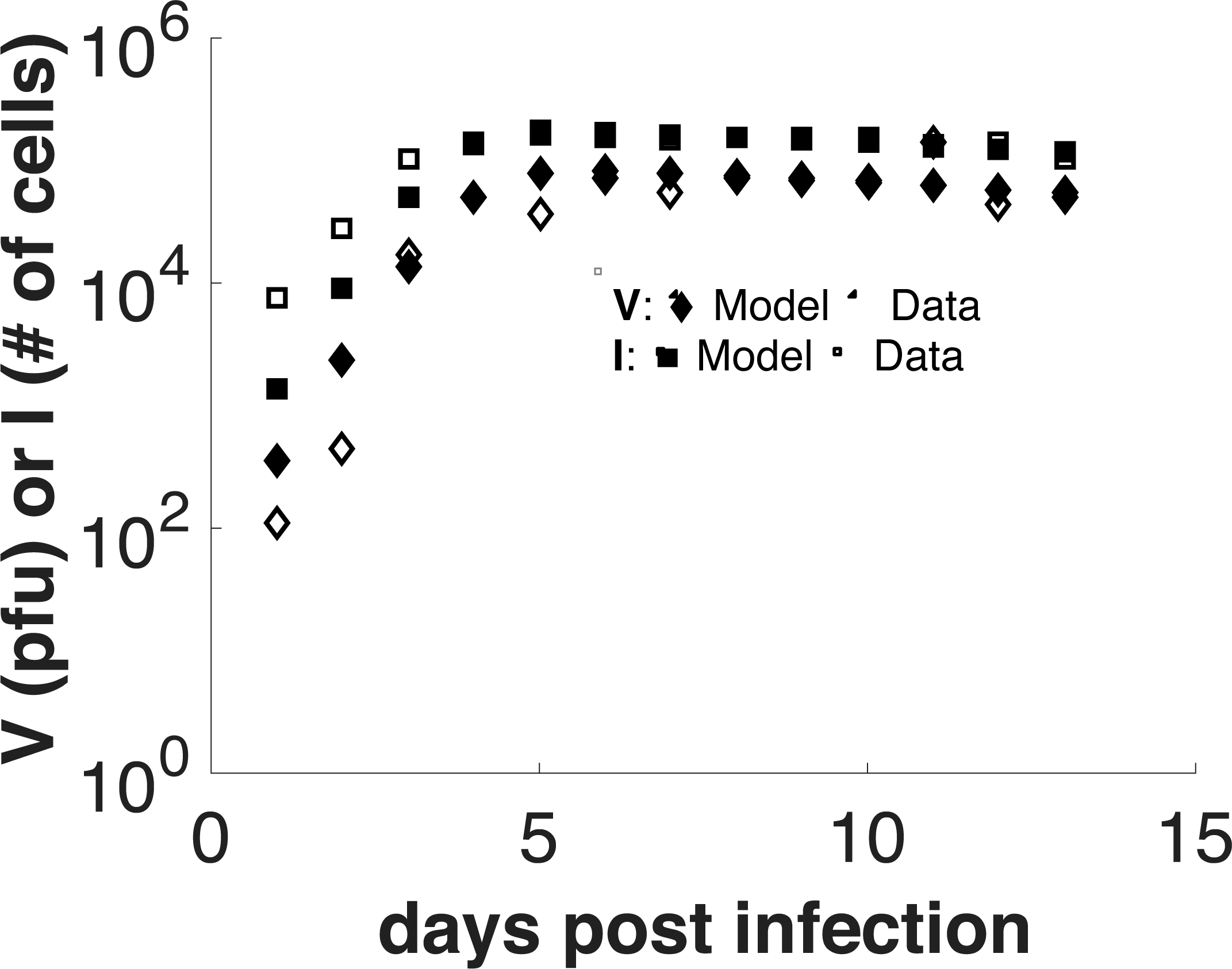
Model fits for the RSV infection *in vitro*. Shows the kinetics of numbers of infected cells (I) and RSV titer (V) measured over 14 days *in vitro*. The model fits are shown with solid symbols. The estimated parameters are listed in Table I. Details of the ODE model are provided in the main text.

**Table I:** Estimated parameter values and confidence intervals (95% CI) for the in vitro experiments.

| $T_0$<br>(# of cells) | $V_0$<br>(# of pfu) | $I_0$<br>(# of cells) | $\beta$<br>(pfu<sup>-1</sup>day<sup>-1</sup>) | $\delta$<br>(day<sup>-1</sup>) | $p$<br>(pfu · cell<sup>-1</sup> day<sup>-1</sup>) | $c$<br>(day<sup>-1</sup>) | $R_0$<br>$\left( \frac{p\beta T_0}{c\delta} \right)$ |
| --- | --- | --- | --- | --- | --- | --- | --- |
| 198,000 | 400 | 0 | $1.317 \times 10^{-5}$<br>($1.087 \times 10^{-5} - 1.546 \times 10^{-5}$) | 0.0555<br>(0.0028 – 0.10814) | 3.83 | 2.3 | 78.24 |

### Estimation of model parameters for RSV kinetics in the lungs of cotton rats

We modeled RSV infection of target epithelial cells (T_0_) present in a gram of lung tissue in cotton rats. We used the data published in Prince et al.^12^ where cotton rats of different ages were infected with the Long strain of RSV. The parameter estimation and their dependence on the age of the cotton rat cohorts were carried out using a non-linear mixed effects modeling approach^16,24^ (details in Materials and Methods section). T_0_ and I_0_ were held fixed at 1×10^7^ cells and 0 for all ages. Assuming 10% of the cells in the lung are ciliated epithelial cells and there are about 10^8^ cells in 1 gram of lung tissue in cotton rats we estimated T_0_ ≈ 1× 10^7^ cells. We first considered *β_a_*,*δ_a_*, 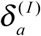, *t_A_*(*a*), *p_a_,c_a_*, and V_0_(*a*) to be random unknown parameters that varied with age *a*. The logarithms of the parameters 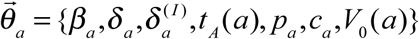 were assumed to vary randomly across ages with a normal distribution with mean values 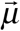 and standard deviations 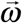^16,30,31^. The differences between the logarithms of viral titers measured *in vivo* (or ν^(data)^) and computed in the model (or ν^(model)^) were assumed to follow a normal distribution with zero mean and a variance σ^2^. The likelihood function 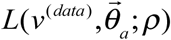, where, 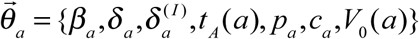 and 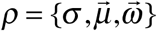, was maximized to estimate the parameter values. Details of the method are described in the Materials and the Methods section. The numerical calculations were performed using the software package Monolix^30^. In order to determine the variability of the maximum likelihood estimate (MLE) of 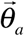 across ages we evaluated the standard deviations 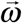 for the parameters (Table S3) which quantified the variations of the parameters across ages. *ω_δ_*(∼ 2.05) and *ω_p_* (∼0.03) showed the largest and the lowest value, respectively. The standard deviations for δ, V_0_, c, and δ^(I)^ were greater than 0.1 and while the rest of the parameters were distributed with standard deviations less than 0.1. The low variability in the parameter values indicated that the age specific variations in the parameters with low standard deviations cannot be resolved well using the available viral titer data points. Therefore, we considered *t_A_* and *p*as age independent but unknown population parameters in our next parameter estimation step. In addition, the estimated value of δ (Table S3) displayed a very small value (∼2.3 × 10^−7^ days ^−1^), thus its effect on the RSV titer kinetics over the duration (∼8 days) of the infection was negligible and we decided to set it to zero in our parameter estimations for the *in vivo* model. The age dependent random parameters were chosen to be 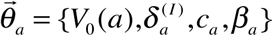 and T_0_ and I_0_ were fixed to the same values as in the previous estimation. The estimated parameters are listed in Table II. The fits to the data are shown in Fig. 5.

**Table IIA:**
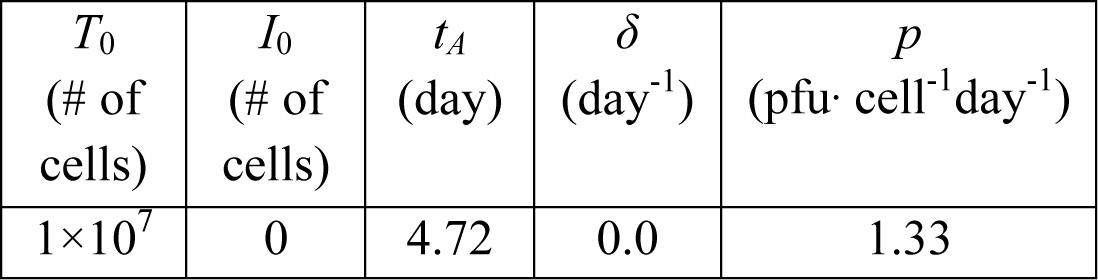
Estimated values for the age independent parameters in the *in vivo* model describing the RSV titer kinetics in the lungs of cotton rats measured in Prince et al.^12^.

| $T_0$<br>(# of cells) | $I_0$<br>(# of cells) | $t_A$<br>(day) | $\delta$<br>(day<sup>-1</sup>) | $p$<br>(pfu· cell<sup>-1</sup>day<sup>-1</sup>) |
| --- | --- | --- | --- | --- |
| $1 \times 10^7$ | 0 | 4.72 | 0.0 | 1.33 |

**Table IIB:**
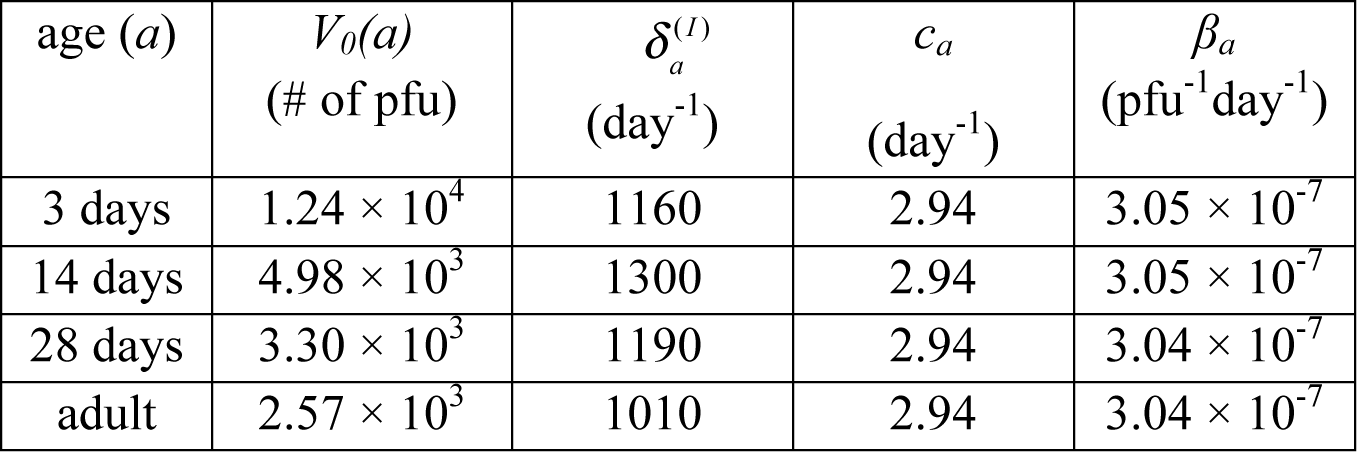
Estimated age dependent parameters V_0_(a), 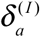, *c_a_*, and *β_a_* for the *in vivo* model describing the RSV titer kinetics the lungs in cotton rats measured in Prince et al.^12^.

**Fig. 5.**
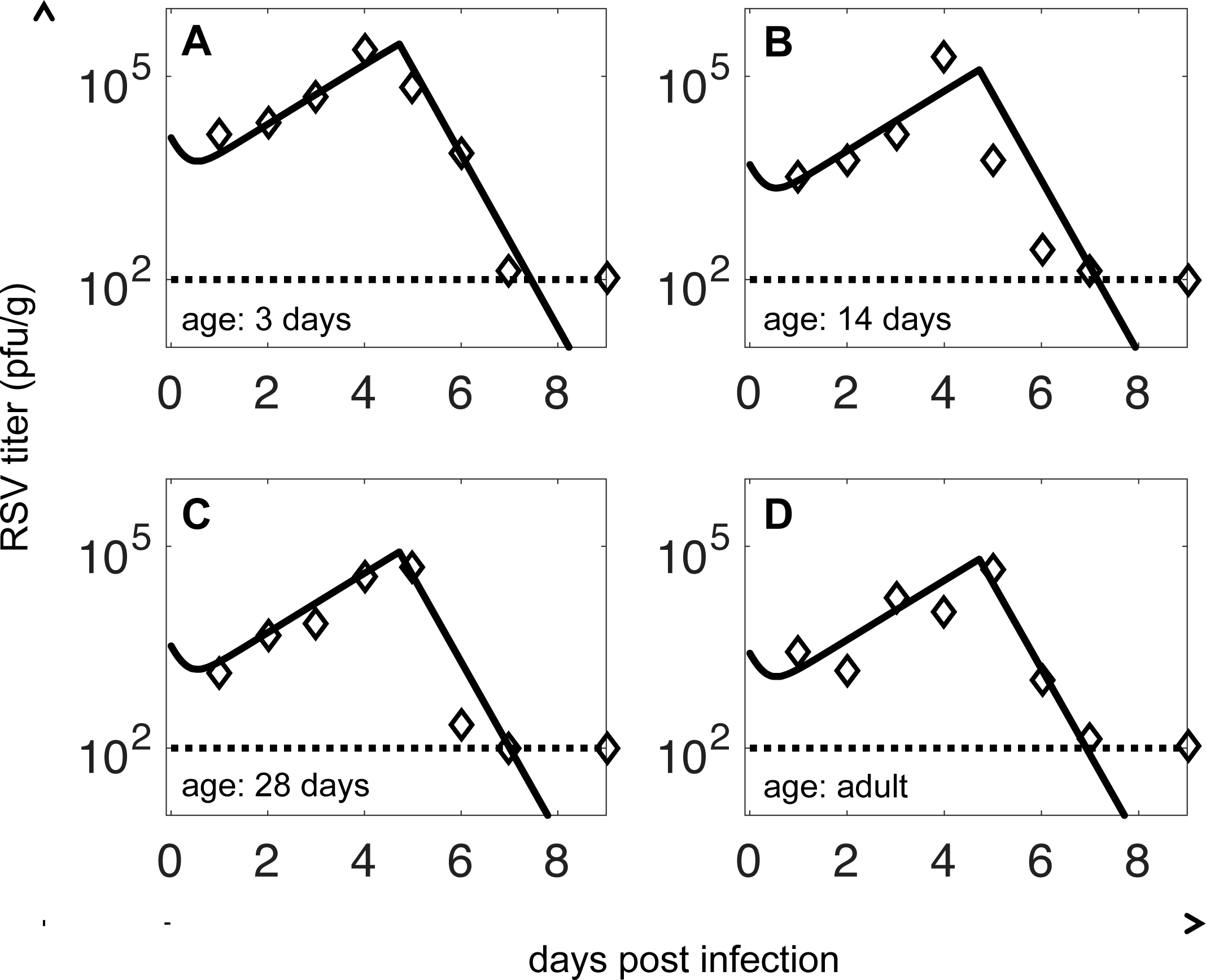
Model fits for the RSV titer kinetics in the lungs of cotton rats. The model fits to the data obtained from Prince et al. are shown for the RSV titer kinetics in the lungs of the cotton rats where the animal cohorts of age (A) 3 days, (B) 14 days, (C) 28 days and (D) 6-8 weeks (adults) were infected with the Long strain of RSV. The estimated parameter values are shown in Table II.

The standard errors, the percentage errors, and the standard deviations 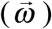 for the distributions pertaining to these parameters are shown on Table S4. The estimates show that V_0_(*a*) to be varying substantially across the age *a* more than any of the other age dependent parameters. V_0_(*a*) also shows a systematic decrease with increasing age which could be due to the increasing size of the lungs as the animal ages (see Discussion). The values of *p* and *c* are in the same orders of magnitudes compared to their values estimated *in vitro*. The decrease in the value of β for the *in vivo* model compared to the *in vitro* model is mainly due to the change in the unit from the *in vitro* to the *in vivo* estimate (see Supplement), and the estimated value of δ or its value for the *in vitro* estimate is too small to have an appreciable effect on the RSV titer kinetics for the duration of the infection. Thus, the RSV titer kinetics *in vivo* appear to be mainly controlled by the introduction of the CD8+ T cell response (non-zero δ^(I)^) in the *in vitro* model.

### Test of model predictions: CD8^+^ T cell response induces faster decline in RSV kinetics

We decreased the value of δ^(I)^ in the best fit model developed in the last section to predict the changes in the RSV titer kinetics when the CD8+ T cell response is suppressed. Our model simulations showed that reducing δ^(I)^ resulted in slower decay of the viral titer after the titer peak (V_peak_) is reached (Figs. 6-7). However, the kinetics before attaining the peak titer remained the same (Figs. 6-7). This prediction was in excellent agreement with the RSV titer kinetics measured in the lungs of the wild-type and the CD8^+^ T cell depleted geriatric (aged over 9 months) (Fig. 6) or adult cotton rats (Fig. 7). We provide further details below regarding how the above prediction was tested against experiments. Cotton rats at different ages, namely, neonates, adult, and geriatric (over 9 months), were inoculated with strain RSV-A2. Cotton rats were inoculated intranasally with 10^5^ TCID_50_ RSV and homogenates from nose and lung were titrated at various days after infection. The non-linear mixed effects model developed for estimating parameters in the previous section was applied to the RSV titers measured in the lungs of the cotton rats at different ages (Fig. S4). The estimated parameters are listed in Table S5. The main differences in the estimated parameters for the Prince et al.^12^ data set and our dataset showed up in *V_0_*(*a*), *p*, *c_a_*, and *δ_a_*^(*I*)^. The estimated values for *p* and *c_a_* are slightly higher than that estimated for the data in Prince et al.^12^, whereas, *V_0_*(*a*) and *δ*_*a*_^(*I*)^ estimations produced lower values compared to that in Prince et al. ^12^. However, the parameter estimations for this dataset also contained larger uncertainties (Table S5) compared to that obtained for the data in Prince et al. ^12^. This is caused by the fewer number of time measurements in this data set. These predictions were compared against RSV titer kinetics measured over 10 days in lungs of CD8^+^ T cell depleted geriatric (Fig. 6) and adult (Fig. 7) cotton rats. The δ^(I)^ values in the best fit models for the adult and the geriatric cotton rats were reduced about 2 fold to generate predictions for RSV titer kinetics in those animals. The model prediction was also tested in cotton rats injected with the drug cyclophosphamide, a drug that suppresses the adaptive immune response by inhibiting cell proliferation. The effect of cyclophosphamide was modeled by decreasing δ^(I)^ tenfold. The RSV titer kinetics in the lungs of cyclophosphamide injected adult cotton rats agreed well with the model prediction (Fig. 7) where the RSV titer kinetics in the cyclophosphamide treated cotton rats did not change appreciably from that of the control during the early stages (<4 days) of the infection but increased substantially at later days (>4 days). The RSV titers measured in the nose of CD8+ T cell depleted or cyclophosphamide treated cotton rats agreed qualitatively with the model predictions as well (Figs. S5-S6).

**Fig. 6.**
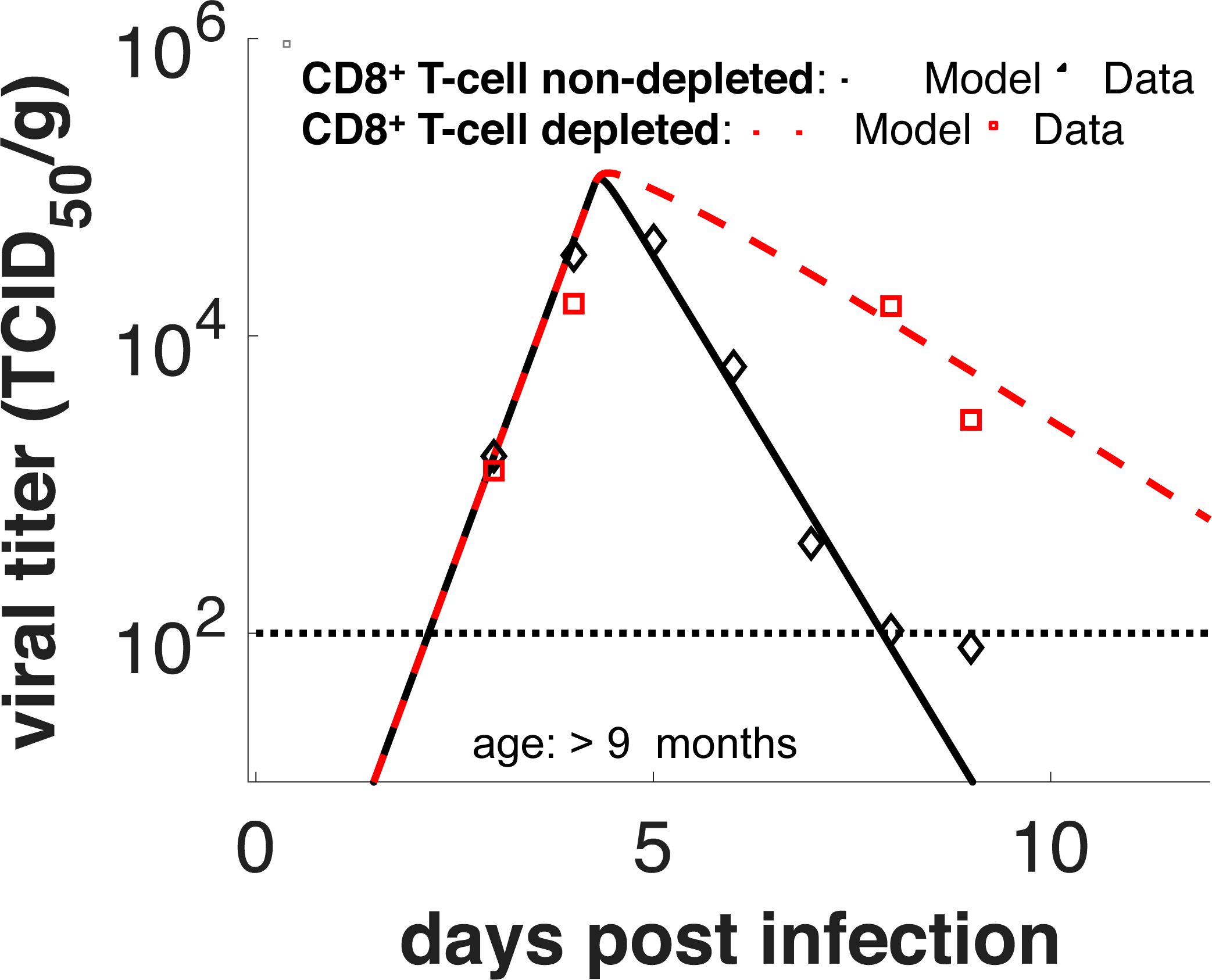
Test of model prediction for the RSV titer kinetics in the lungs of CD8 T cell depleted geriatric cotton rats. The model predicted (grey and red dashed lines) that reduction of the strength (δ^(I)^) of the CD8 T cell response by approximately 2 fold does not affect the RSV titer at early stages (<4 days) of infection, however, it increases the RSV titer at later days (> 4 days). This model prediction was tested by measuring RSV titer kinetics in the lungs of the CD8 T cell non-depleted and CD8 T cell depleted geriatric (age ∼ 270+ days) cotton rats. As predicted by the model, the RSV titers in the CD8 depleted cotton rats do not change appreciably from their counterpart in the CD8 T cell non-depleted at 3 and 4 dpi, however, at later stages of infection (5 and 6 dpi), the RSV titers in the lungs of the immunosuppressed cotton rats are substantially larger than that in the CD8 T cell non-depleted cotton rat.

**Fig. 7.**
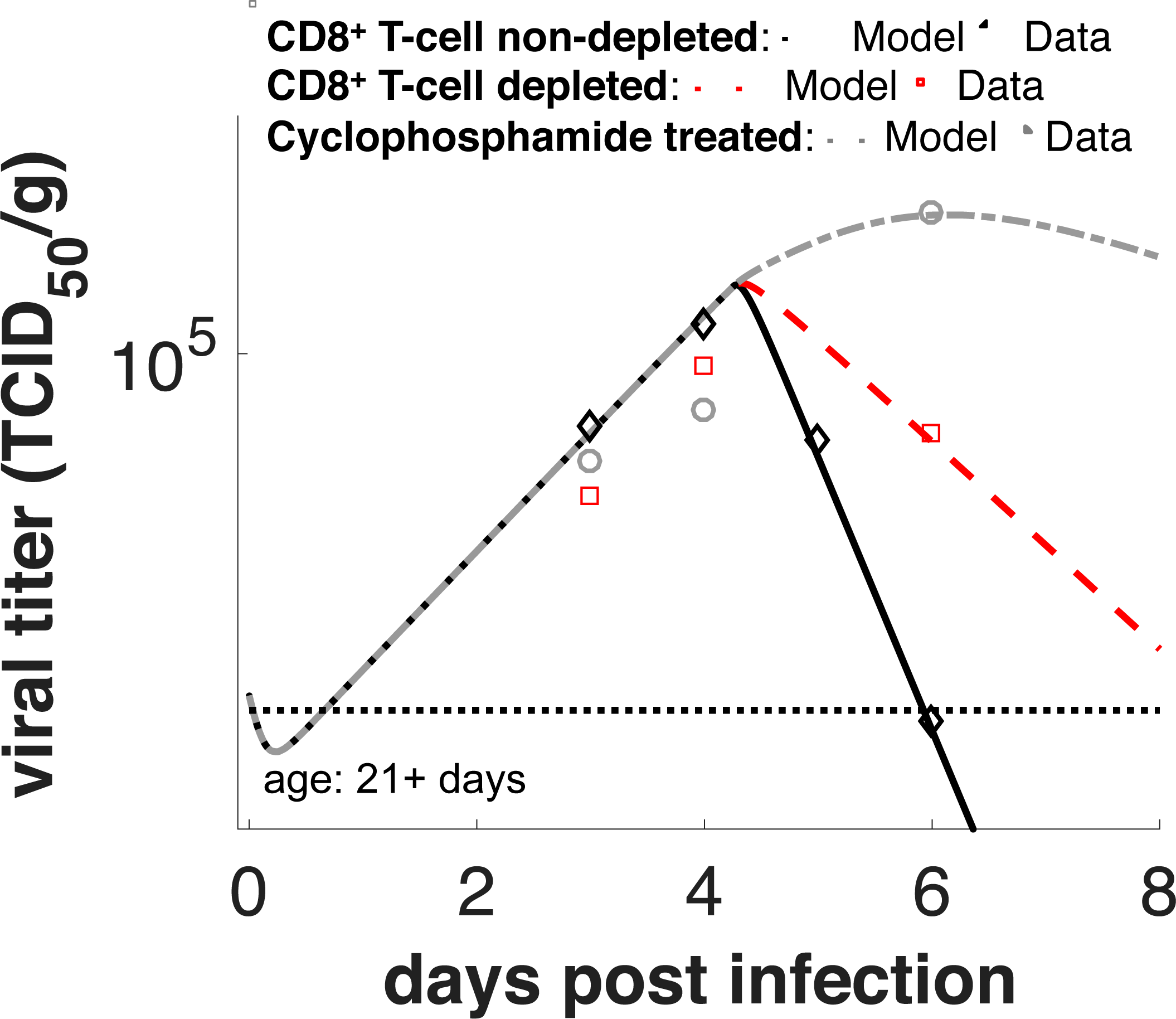
Test of model prediction for the RSV titer kinetics in the lungs of CD8 T cell depleted or cyclophosphamide treated adult cotton rats. The model prediction regarding the reduction of CD8 T cell response was tested by measuring RSV titer kinetics in the lungs of CD8 T cell non-depleted, CD8 T cell depleted, and cyclophosphamide treated geriatric (age ∼ 270+ days) cotton rats. In cyclophosphamide treated cotton rats all rapidly replicating cells are depleted thus the adaptive immune response is severely blocked in these animals. Depletion was modeled by reducing δ^(I)^ by approximately 2-fold and cyclophosphamide treatment was modeled by reducing δ^(I)^ by 10 fold. The varying effects only take place after ∼5 days. Similar to Fig. 7, the model prediction is in excellent agreement with the measurements.

### Discussion

We developed a simple mathematical model using ODEs to describe RSV titer kinetics in the lungs of cotton rats. The model was able to describe several unique features of RSV titer kinetics and demonstrated the importance of the CD8+ T cell response in the decline of the RSV titer kinetics at later stages (> 4 days) in the lungs of cotton rats. The ODE model generated quantitative predictions regarding the RSV titer kinetics in CD8+ T cell depleted or immune-suppressed cotton rats which were in excellent agreement with experiments. Therefore, we have developed an experimentally validated quantitative model with predictive powers for describing RSV titer kinetics in cotton rats.

The ODE model developed for RSV titer kinetics *in vivo* was modified to describe RSV titer kinetics *in vitro* where HAE cells were infected with a human strain of recombinant RSV expressing green fluorescent protein (rgRSV). The model incorporated specific steps following the *in vitro* experiments where RSV virions were washed from the culture wells every 24 hours and measured (details in Materials and Methods section). We estimated model parameters for the RSV kinetics by fitting the model with the measured values of RSV titers in the supernatant and the counts of infected cells. The estimation of the reproductive number^29^ R_0_ *in vitro* is ∼80 suggesting a high infection capability (i.e., ∼80 new infections initiated on average by a single infected cell during its lifetime) of RSV. The estimated replication rate *in vitro* is high, (∼3 pfu per day per infected cell), and the average time scale for a single HAE cell death due to RSV infection is ∼ 50 days, which indicates that RSV infection is not readily cytopathic. The low rate of infected cell death and large RSV replication rate contributed towards the large value of R_0_ *in vitro*.

The parameter estimation for the *in vivo* data were much less constrained compared to the *in vitro* data as the RSV titers alone were measured. Non-linear mixed effects modeling^16,24^ estimated the parameters, as well as age dependence of some of the parameters that captured variations of the RSV titer kinetics with the age of the cotton rats. Most of the parameter estimates (β, p, δ, c) for RSV kinetics in the lung measured by Prince et al. ^12^ were similar to that estimated for the *in vitro* measurements and these parameters did not vary substantially (standard deviation/(mean)<0.1) with the age of the cotton rats. The animals were infected with the Long strain of RSV in the studies by Prince et al. ^12^. The age dependency of the RSV titer kinetics in the lung appeared to be predominately captured by the systematic decrease of V_0_with increasing age. Since the size of the lung increases as cotton rats age, the amount of inoculum initiating infection in a fixed volume of lung (or V_0_) decreases with increasing age. This could underlie the age dependent behavior of V_0_ we found in our parameter estimation. δ^(I)^ varied randomly across age by a very small amount (<2%) and thus its role across age appears to be negligible.

The values of model parameters appeared to depend on the strain of the RSV used in infecting the cotton rats. Some of the estimated parameters (*p* and *c*) for the RSV kinetics measured in our experiments where cotton rats were infected by RSV-A2 strain were slightly higher compared to that estimated for the data in Prince et al. The age dependency of V_0_ did not show a systematic change with age and c showed larger variations with age (Table S5). However, the uncertainties in the parameter estimates were also higher for our dataset due to measurements at fewer time points. This is a common scenario in estimation of parameters of non-linear ODEs where only a subset of the variables is measured^20,32^. However, it is still possible to generate quantitative and discernable predictions by perturbing certain parameters^33^. In our case, we decreased δ^(I)^ to generate predictions of RSV kinetics in CD8+ T cell depleted or immunocompromised cotton rats in excellent agreement with the experiments. This showed the utility of generating falsifiable predictions in situations where several model parameters cannot be well estimated.

We characterized the geometry of the RSV titer kinetics using three features that impose constraints on mechanisms that can be used in constructing ODEs for describing viral titer kinetics. A simple mathematical model^14^ for IAV infection was unable to produce viral titer kinetics with τ_ratio_<1 for a wide range of parameters. In order to generate viral titer trajectories with τ_ratio_<1 one needs to include a term in the ODE model for IAV infection that comes into action after a time lag, which in our case represented the clearance of RSV infected cells by CD8+ T cells. This effect was implanted minimally by using a step function in our ODEs, however, a more realistic way to include this effect could be to model many intermediate activation steps with different time scales which can potentially give rise to such time lags^34^.

Several RSV titer mean trajectories in the nose and trachea of cotton rats contained more than one peak. However, it is difficult to assess if the RSV titer in the same animal showed multiple peaks over time because the animals were sacrificed at each measurement. Given the small sample size of the animal cohort (3-4 cotton rats per cohort) the presence of the multiple peaks in the geometric average of the RSV titer could be attributed to animal-to-animal variation of the RSV titer. However, if the multiple peaks in the geometric average of the RSV titer are representative of the actual RSV titer kinetics in an individual cotton rat, such behavior can arise due to the interplay between induction of interferons (e.g., IFNγ) by RSV and the suppression of viral replication by the interferons. Baccam and colleagues^14^ showed a time delay in the induction of the interferons and the suppression of the viral replication by the interferons could produce multiple peaks in IAV titer kinetics. This is because the interferons decrease viral load by suppressing viral replication, however, lowering the viral load also decreases the induction of interferons thereby increasing viral replication, thus, the viral titer decreases after the first peak and then increases again to generate a second peak. Similar mechanisms can be included in the simple model to describe potential multiple peaks in the RSV titer as well.

The approach carried out here combining simple mechanistic modeling and measurements of RSV titers shows the usefulness of simple mathematical models in deciphering mechanisms that underlie RSV production kinetics in cotton rats. The model developed here can be used to assess outcomes of vaccine candidates or anti-viral therapies against RSV in cotton rats or other animal models.

### Materials and Methods

#### Non-linear mixed effects modeling of RSV titer kinetics in cotton rats of different ages

The temporal profiles of RSV titers measured in the lungs of the cotton rats showed differences with respect to age(Fig. 3A). Therefore we modeled the RSV infection kinetics in cotton rats with parameter values that depended on the age (*a* in {3 days,14 days, 28 days, Adult}) of the animal. We employed a non-linear mixed effects approach that models age-specific parameters as random effects. For each time point *t* in (1, 2, 3, 4, 5, 6, 7, 9, 12, 15, and, 20) days post inoculation and for each age *a*, we computed the logarithm of the geometric mean RSV titer (denoted log_10_ V^(data)^(t,a)) at a particular time *t*, where the geometric mean was taken over 3 cotton rats in an animal cohort of age*a*. The parameters of the non-linear mixed effects model were estimated by maximum likelihood (ML), which (provided that there is a unique maximum) finds the set of parameter values that makes the RSV titer log_10_[V^(data)^(t,a)] most probable. Given the age-specific random effects parameters 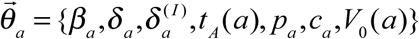, and the fixed parameters (T_0_ and I_0_), 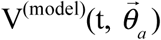 was obtained by numerically solving the ODEs in Eqs. (1)-(3). For brevity we will denote log_10_[V^(data)^(t, a)] and 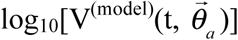 and 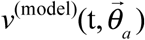, so that the non-linear mixed effects model is more succinctly expressed by: 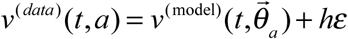. Here, the residual error (denoted *hε*) may depend on fixed parameters (or measured covariates) through the function *h*, whereas, *ε* is distributed as a normal distribution 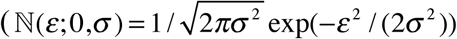) of zero mean and an unknown variance *σ*. We assume that *ε*(*t*) is independent of age and *h*=1. We also fixed σ to the standard deviation in the dataset σ^(data)^ = 1/(# of timepoints × # of age groups × # of animals in the animal cohort)∑_{t,a,α}_ (ν^(data)^_(α)_(t,a) - ν^(data)^ (t,a))^2^ ≈ 0.5, where ν^(data)^_(α)_(t,a) denotes the log of the RSV titer measured at time *t* in a cotton rat (indexed by α) that belong in the animal cohort of age *a*. For our dataset for inoculation with RSV-A2 strain σ^(data)^=0.23. 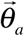 is given by the sum of two components: unknown but fixed (i.e., age independent) parameters 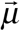, and, random parameters 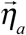 that depend on the age *a*, *i.e.*, 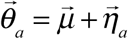. The random vector 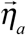 is assumed to be independent and identically distributed (iid) following a specified probability distribution function (e.g., normal distribution with zero mean and an unknown variance 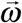). The joint probability distribution function is given by 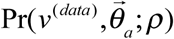, where, 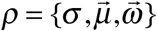, is the likelihood function: 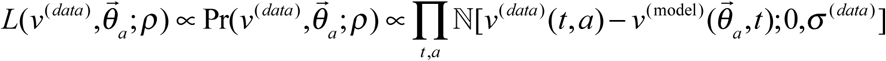. The likelihood function 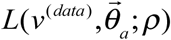 is maximized to evaluate the maximum likelihood estimates (MLE) of the unknown fixed parameters, namely, σ and 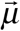, and, random parameters 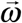.

The probability distribution for each age-specific parameter vector 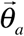 is estimated from the conditional probability distribution, 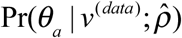, where 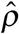 is the ML estimate of ρ. Computing the ML estimates of the model parameters involves optimization of high-dimensional cost functions. The landscape of the cost function for non-linear models is typically multimodal, containing more than one local maximum. As such, parameter estimates are often dependent on the initial guesses for the parameter values, which are then used to seed the search. In order to get a sense of the range and the initial guess for the model parameters we first fitted the model for measurements in an *in vitro* experiment where both the RSV load and number of infected cells can be measured at many time points and the inoculation dose and the number of target cells can be clearly controlled. The *in vivo*estimations were carried out using the *in vitro* estimates of the common parameters (Table I) as initial conditions. The unit of β was changed for the *in vivo* models following the formula derived in the Supplemental Text. In the Monolix software package, we used decreasing variability to zero for fixed parameters, with 5 burn-in iterations, 10,000 exploration iterations, 200 smoothing iterations (no auto-stop permitted), and the simulated annealing option on.

#### In vitro measurements

##### Preparation of HAE cultures

###### HAE cell recovery and culture

The nasal respiratory epithelium was sampled using a cytology brush and the cells were recovered as previously reported ^35^. Mucus was removed through treatment with 14.3 µM 2-mercaptoethanol and cells were cultured using the mCRC method ^35^. At passage 1, all cells in the culture were basal cells. Basal cells were plated onto collagen-coated 0.33 cm^2^ transwell membranes at 2X10^4^ cells per membrane as previously described ^35^. At confluence, the medium was changed to PneumaCult-ALI Base Medium containing the 10X supplement, the 100X supplement, heparin, and hydrocortisone (Stem Cell Technologies, Vancouver, BC, Canada).

###### Inoculation of HAE cultures with RSV, imaging, and virus yield

On differentiation day 21, HAE cultures were washed with DMEM to remove mucous and inoculated with 400 plaque forming units (pfu) of recombinant green fluorescent protein (GFP) expressing RSV strain A2 (rgRSV224) ^36^ in 50 µL of DMEM for 4 hr at 37°C. Each day post-infection wells were imaged to quantify GFP fluorescence. Fluorescent was quantified with the MIPAR program (MIPAR Image Analysis; Worthington, OH) by multiplying the mean intensity fluorescence by the surface area fraction of fluorescence. A separate set of wells were washed with 100 µL of DMEM containing 10% FBS, and the wash was snap frozen on dry ice and stored at −80°C for later quantification. The number of infectious viruses in each wash was quantified by serial dilution titration on HEp-2 cells in 96-well plates and counting the green fluorescent foci at 24 hr post-inoculation.

##### In vivo measurements

###### Animals

Inbred cotton rats (*Sigmodon hispidus*) were obtained from Envigo, Inc. (Indianapolis, IN) and housed in standard polycarbonate cages in a barrier system, or obtained from an in house breeding colony. Corncob bedding (Cincinnati Labs Supply) was used for all experimental animals. Environmental conditions were constant at 22±2 °C, 30-70% relative humidity, and 12:12 hours light:dark cycle. Cotton rats of both sexes between day 3 and 8 weeks of age were used. Euthanasia was performed via CO_2_ inhalation. All animal experiments were approved by the Institutional Animal Care and Use Committee of The Ohio State University. Cotton rats were purchased specific pathogen-free and maintained in colonies free of endo- and ectoparasites, mouse parvovirus 1 and 2, minute virus of mice, mouse hepatitis virus, murine norovirus, Theiler murine encephalomyelitis virus, mouse rotavirus, Sendai virus, pneumonia virus of mice, reovirus, Mycoplasma pulmonis, lymphocytic choriomenginitis virus, mouse adenovirus, and ectromelia virus according to quarterly health monitoring of sentinel (CD1) mice exposed to 100% pooled dirty bedding from colony animals at each cage change.

###### Infection

Cotton rats were anesthetized via isoflurane inhalation before being inoculated intranasally with virus diluted in PBS to a final inoculation volume of 100 µl (10ul for neonates).

###### Virus

Stocks of RSV A/2 were grown in HEp-2 cells in MEM/2% fetal bovine serum. When infection reached a cytopathic effect of ∼80%, cells were scraped from the flask in the growth media. The cells were briefly frozen at −80°C, thawed, and centrifuged at 3 000 rpm for 15 minutes at 4° C to remove all large cellular debris. The supernatant was collected and placed on top of 15 ml of a 35% sucrose cushion and centrifuged at 15 000 rpm in a Sorval SS 34 rotor for 5 hours at 4° C to pellet the virus. Virus pellets were re-suspended in MEM/10% trehalose, and TCID_50_ was determined by titration on Hep-2 cells^37^. Each virus stock was aliquoted and frozen in liquid nitrogen. Aliquots were thawed immediately prior to use and were only used once to prevent loss of titer due to repeated cycles of freeze-thaw.

###### CD8^+^ T cell depletion

Cotton rats were inoculated on day −1, 1 and 3 after infection with 0.5 mg of a cotton rat CD8 alpha specific monoclonal antibody^38^.

###### Cyclophosphamide treatment

Cotton rats were injected with 150 mg/kg cyclophosphamide (Sigma-Aldrich) intraperitoneally on day −4, day −2, and day 0 of infection.

###### Statistical analysis

Data are represented as means ± SD. Statistical significance was determined by multiple t tests using the Holm-Sidak method, with alpha = 0.05, and each cell type was analyzed individually, without assuming a consistent SD. Data were analyzed using Prism version 7.00 for Windows, (GraphPad Software, La Jolla, California, USA).

## Acknowledgements

This work was partially supported by grants R01GM103612 to JD and P01 AI112524 to MP and SN, and from the Cystic Fibrosis Foundation to MP.DW and JD thank Alan Perelson for a stimulating discussions.

## Supplementary Material for “Mathematical Modeling Identifies the Role of Adaptive Immunity as a key Controller of Respiratory Syncytial Virus (RSV) Titer in Cotton Rats”

**Figure S1:**
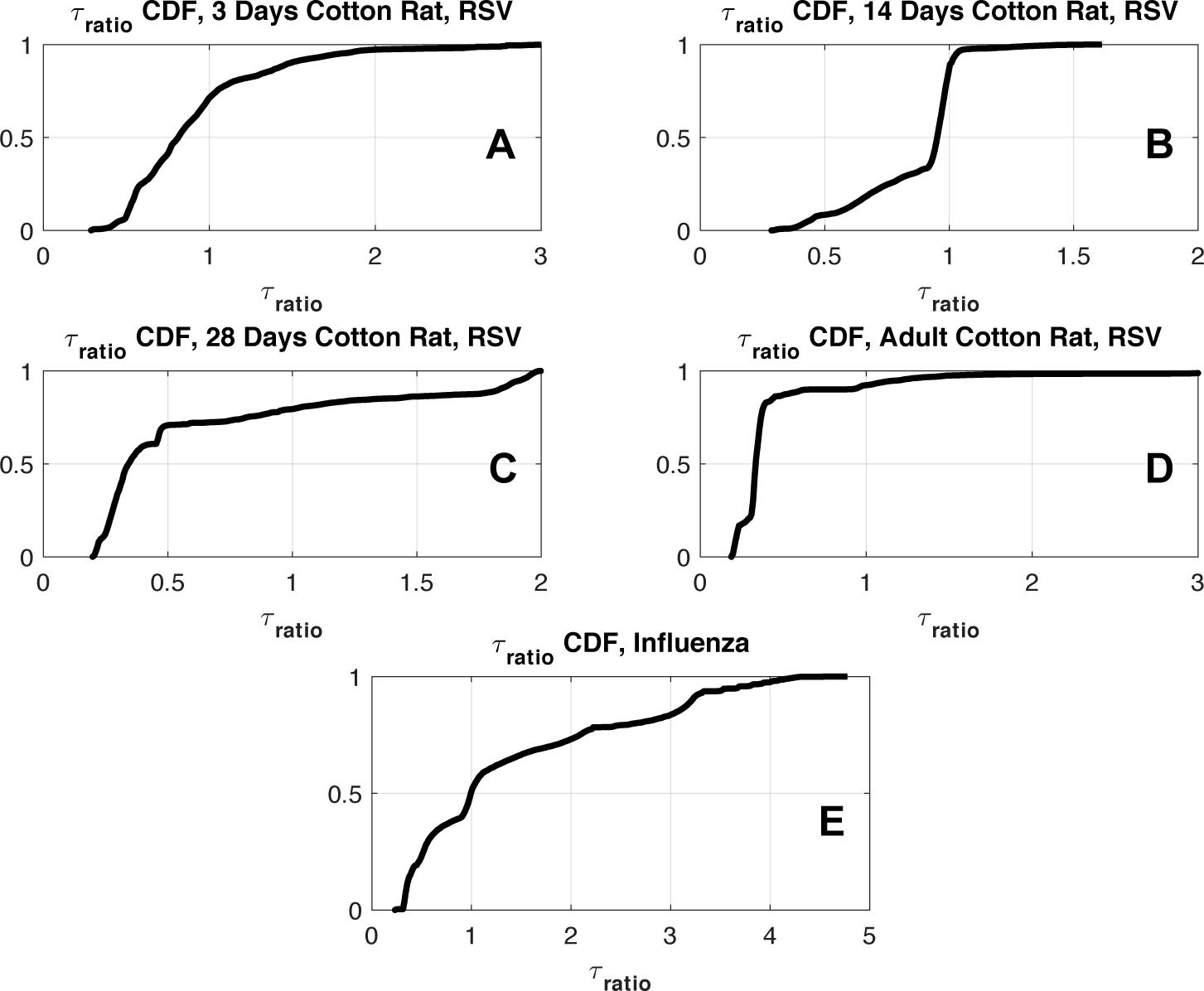
Quantification of uncertainties for estimation of τ_ratio_ for RSV titer kinetics. RSV viral load in Prince et al. was recorded in three cotton rats each day. Thus, the geometric average is likely to be different from the true expectation value due to small number of samples. We estimated the uncertainty in τ_ratio_ by generating potential RSV titers in the animal cohort by bootstrapping where we randomly selected three RSV titers in individual cotton rats from the available values at each day and calculated their geometric means. With 10 possible combinations of data (and thus 10 possible averages) at each of 7 time points, it is possible to create 10^7^ unique trajectories. We calculated τ_ratio_ for these random trajectories and generated probability distribution function P(τ_ratio_) of the possible values. The variation of the cumulative probability distribution function, defined as, 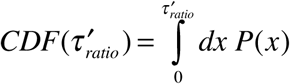 is shown. CDF(τ_ratio_) denotes the probability for a RSV titer kinetics to generate a τ_ratio_ between 0 and τ’ratio. The data are shown for the cotton rat cohorts for ages **(A)** 3 days, **(B)** 14 days, **(C)** 28 days, and **(D)** adult cotton rats. We find that >80% of the trajectories generated from the bootstrapped samples produced τ_ratio_ values less than unity. **(E)** We compared our results against the IAV titer kinetics in human subjects available from Baccam et al.. In contrast to RSV titer kinetics in cotton rats, <50% of the IAV titer kinetics generated τ_ratio_ between 0 and 1. This demonstrates a qualitative difference in the geometric shape between the IAV titer kinetics and RSV kinetics.

**Figure S2:**
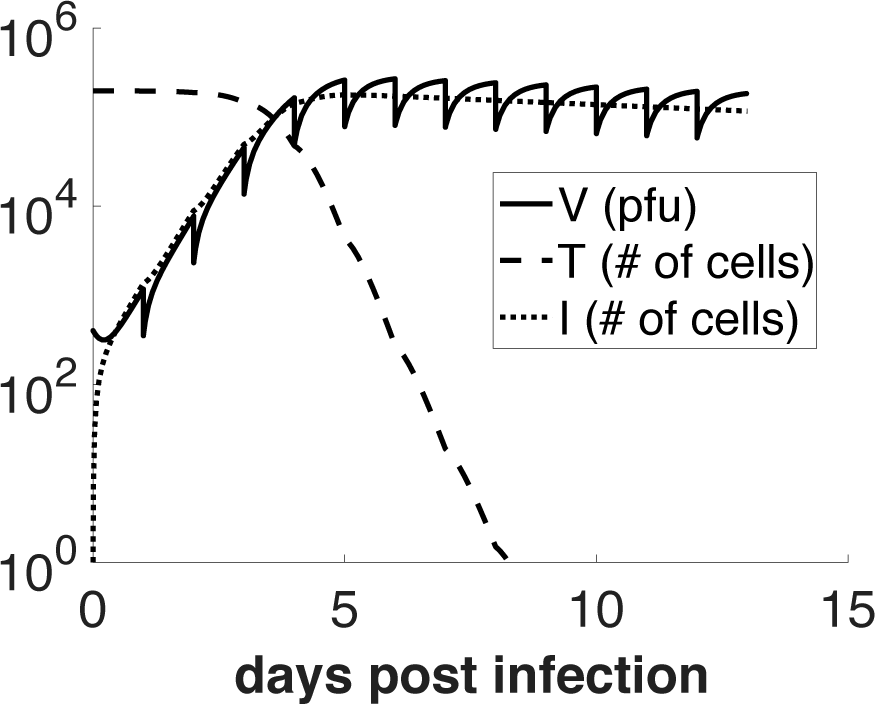
Kinetics of T, I and V in the ODE model describing RSV infection of HBE cells in vitro. Shows the kinetics of the numbers of target cells (T), infected cells (I), and the free RSV virions (V) in the ODE model where the free RSV virions were removed (or washed out) each day for measuring the viral titer. The ODE model is solved following the method described in the main text. The sudden decrease at each day in the viral titer represents the removal of the medium for the measurement. We assumed that a fixed fraction λ of the V is removed during the measurement. λ is set to xx for the data shown above. The other parameters are set to the values in Table I in the main text.

**Figure S3:**
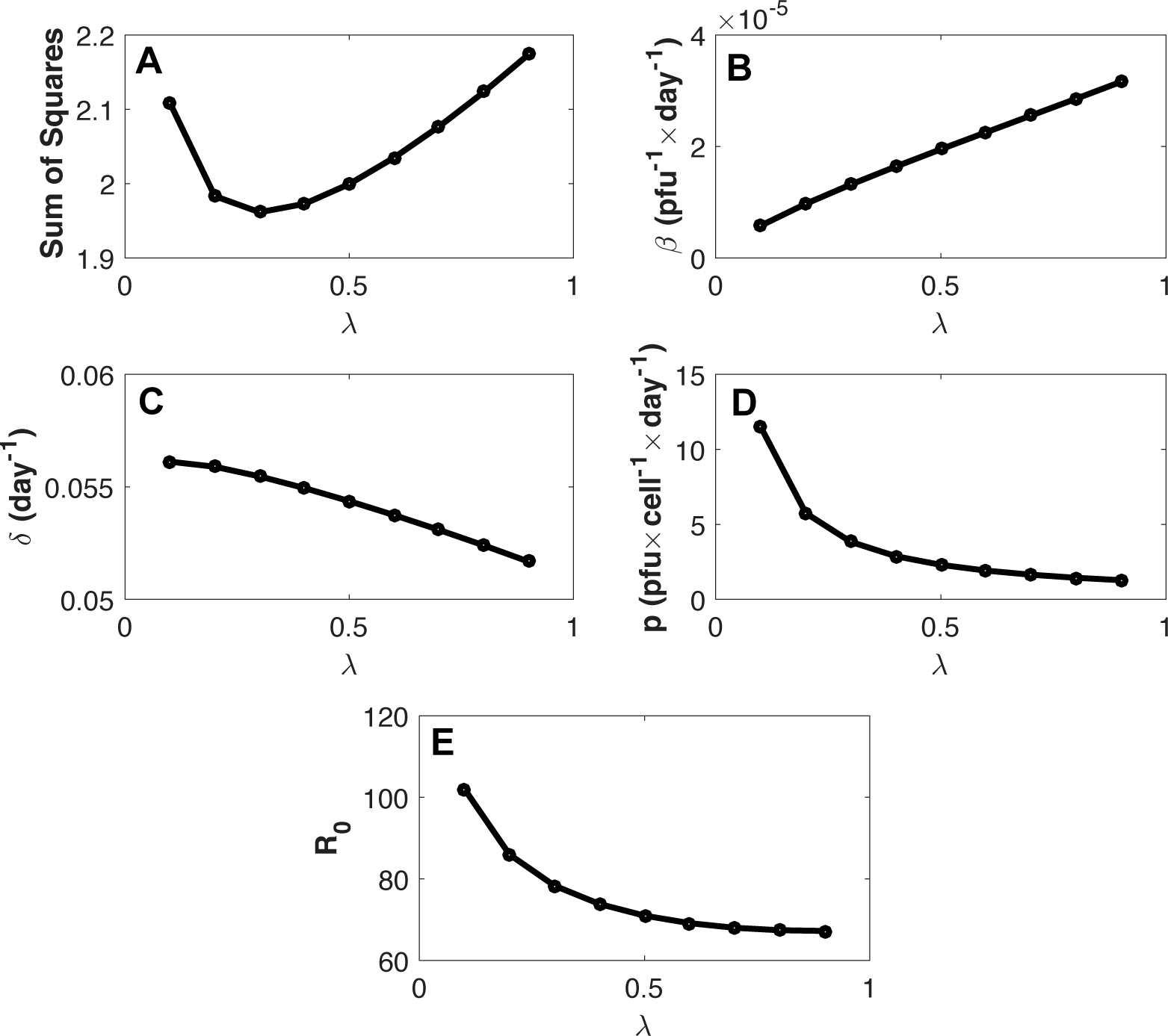
Dependence of the estimated model parameters on the fraction (λ) of virus removed at measurement. Shows the dependencies of the **(A)** sum of least square difference between the model and the data, **(B)** *β*, **(C)** *δ*, **(D)** *p*, and **(E)** *R*_0_ on λ. The least square shows a minimum at λ=0.3. The changes in the estimated parameters *p* and *R*_0_ decrease for λ > 0.5. The changes in δ with increasing λ is very small (∼10%). β increases almost linearly with λ, however, the values reside in the same order of magnitude (O(10^−5^)).

**Figure S4:**
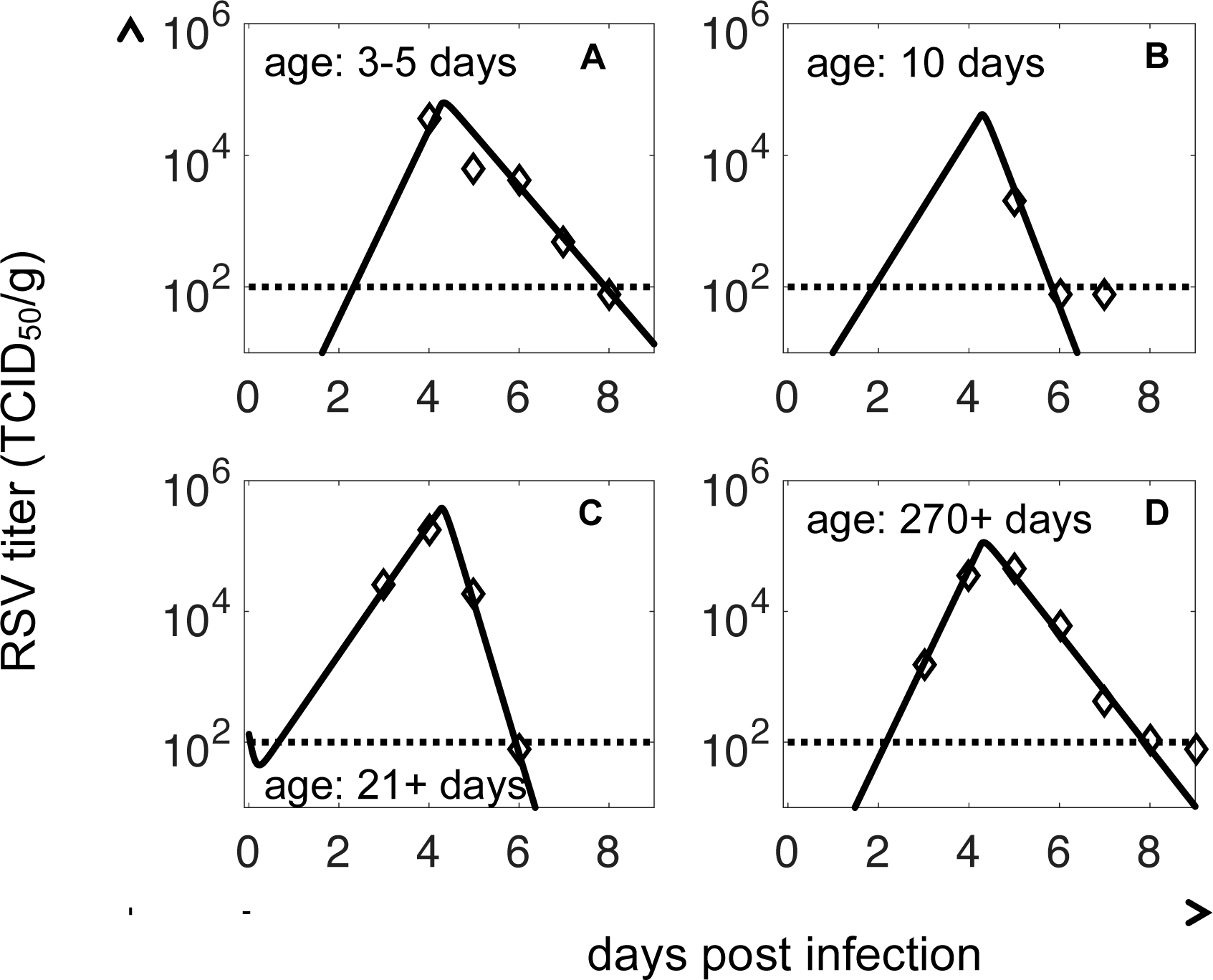
Model fits to RSV titer data in the lungs of cotton rats inoculated by RSV-A2 strain. The fits for cotton rat cohorts of age **(A)** 3-5 days, **(B)** 10 days, **(C)** 21+ days, and **(D)** 270+ days (geriatric) were generated by the nonlinear mixed effects approach described in the main text. The parameter values are shown in Table S5. There appears to be a faster rate of viral clearance after the peak titer is reached in the adult cotton rats as opposed to the neonate or geriatric cotton rats.

**Figure S5:**
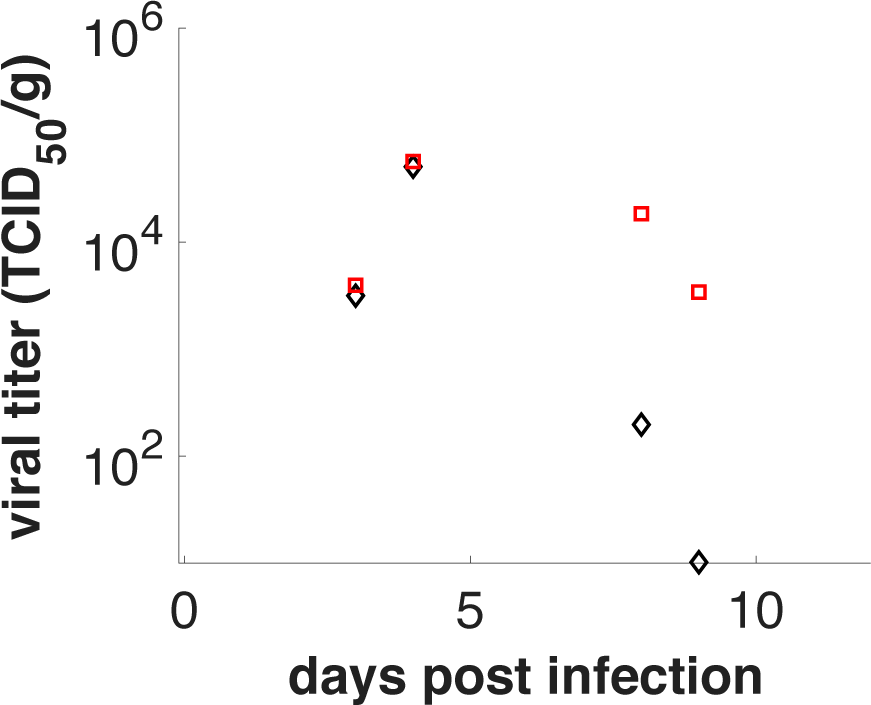
Comparison of RSV titer kinetics in the nose of CD8^+^ T cell depleted and CD8+ T cell non-depleted geriatric cotton rats. Viral titers at early times before the peak titer is reached are similar between CD8 T cell depleted (red squares) and CD8^+^ T cell non-depleted (black diamonds) cotton rats. In contrast, CD8 T cell depleted cotton rats show higher viral titers at later times. This behavior is in excellent agreement with the model prediction. This implies that the CD8+ T cell response plays an important role in controlling RSV titer in the nose of geriatric cotton rats.

**Figure S6:**
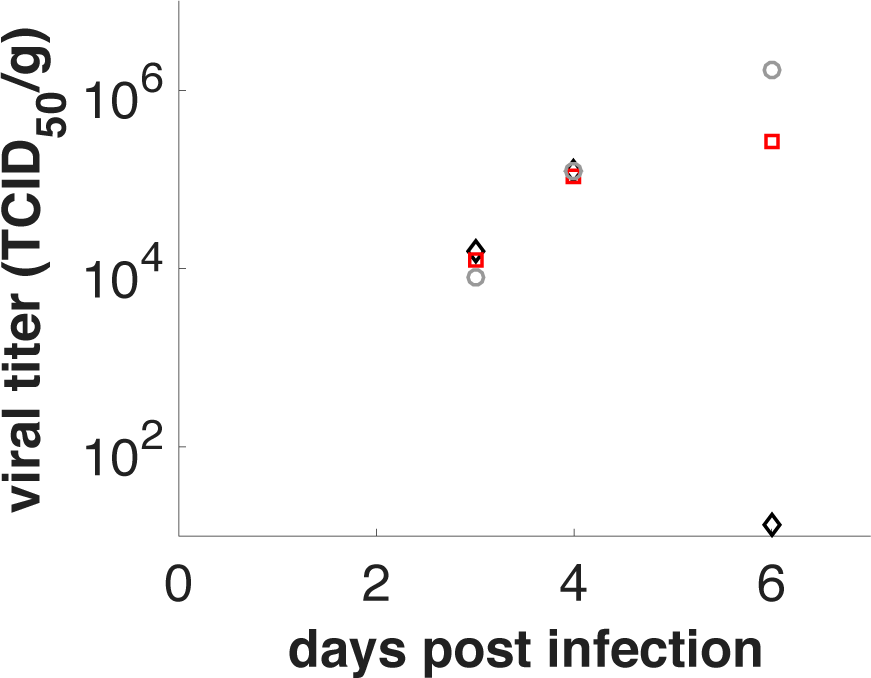
Comparison of RSV titer kinetics in the nose of immunocompromised and non-immunocompromised adult cotton rats. Viral titers at early times before the peak titer is reached are similar between cyclophosphamide treated (grey circles), CD8 T cell depleted (red squares), and non-immunocompromised cotton rats (black diamonds). In contrast, the cyclophosphamide cotton rats show higher viral titers at later times. In cyclophosphamide treated cotton rats all rapidly replicating cells are depleted thus these rats the adaptive immune response is severely blocked in these animals. This behavior is in excellent agreement with the model prediction. This implies that the CD8+ T cell response plays an important role in controlling RSV titer in the nose of adult cotton rats.

**Figure S7:**
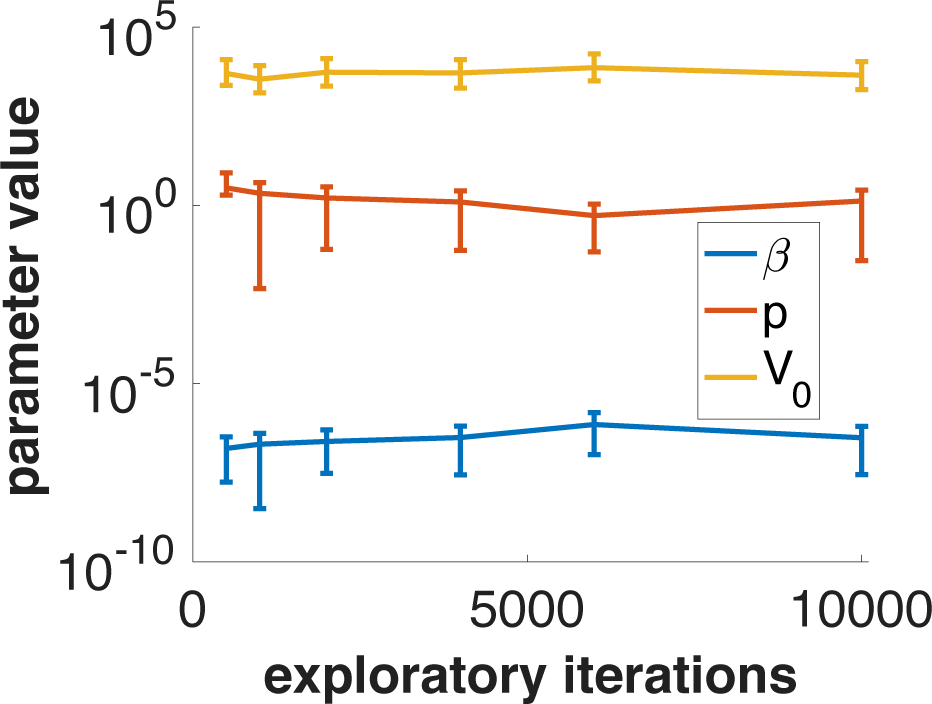
Dependence of Parameter Values on Number of Monolix Iterations. All parameter variations shown are within 1 log. Parameters not showed display similar dynamics. This indicates stability of parameter estimates with respect to the settings of the SAEM algorithm.

## Supplementary Text: Change of units in model parameters

Consider N_T_, N_I_, and, N_V_ number of target cells, infected cells, and virions in a volume of *v*. The densities of these variables are denoted by, T≡N_T_/*v*, I≡N_I_/*v*, and, V≡N_V_/*v*. The viral kinetics described in terms of the density variables is given by the following ODEs,

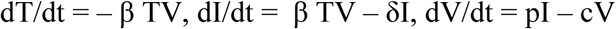

The viral kinetics in terms of the numbers, N_T_, N_I_, and, N_V_ are given by,

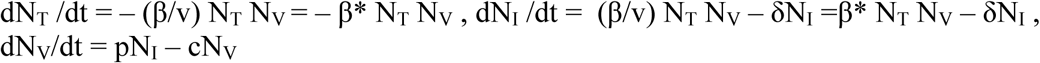

where, β* = β/*v*.

If the same type of target cells, infected cells, and virions interact with each other but in a different volume, *v’* (>*v*), then the densities of these variables (say, T’, I’, and V’) follow the kinetics,

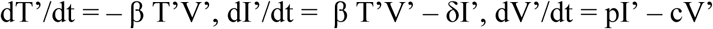

Thus, the corresponding β* for this case (say, (β*)’) will be related to β as

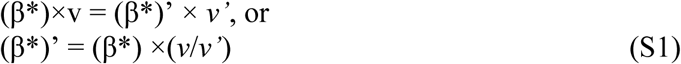

We use Eq. (S1) to change our unit of β* when we change the units from the *in vitro* to the *in vivo* system.

**Table S1:**
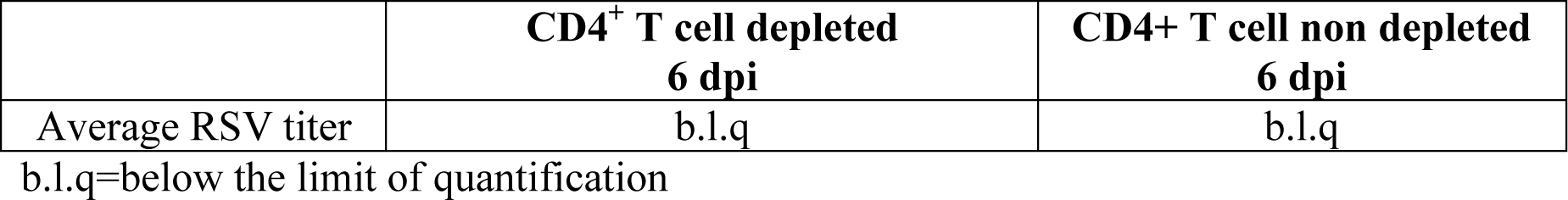
Comparison of RSV infection in CD4+ T cell depleted and CD4^+^ T cell non-depleted cotton rats. Both CD4^+^ T cell depleted adult cotton rats and CD4+ T cell non-depleted cotton rats show full viral clearance by 6 days post infection. Both animal cohorts were inoculated with the same dose (10^5^ TCID_50_). This indicates that CD4^+^ T cells do not play a major role in clearing RSV infection at later days.

**Table S2:** Ranges of parameter values used for generating Fig. 3A.

| | $\beta$ | $\delta$ | $p$ | $c$ | $\delta^{(I)}$ | $t_A$ | $V_0$ |
| --- | --- | --- | --- | --- | --- | --- | --- |
| RSV model<br>(Eqs. (1)-(3)) | $10^{-6}$ to<br>$10^{-10}$ | $2 \times 10^{-4}$ to<br>2 | 0.3 to<br>3000 | 0.0377 to<br>377 | 0.1 to<br>1000 | 0 to<br>15 | 10 to<br>100000 |
| Flu model<br>Baccam et al. | $2.7 \times 10^{-7}$ to<br>0.0027 | 0.04 to<br>400 | $1.2 \times 10^{-4}$ to<br>1.2 | 0.03 to<br>300 | NA | NA | $9.3 \times 10^{-4}$ to<br>9.3 |

$T_0$ was held fixed at $4 \times 10^8$ cells for both the models.

**Table S3:**
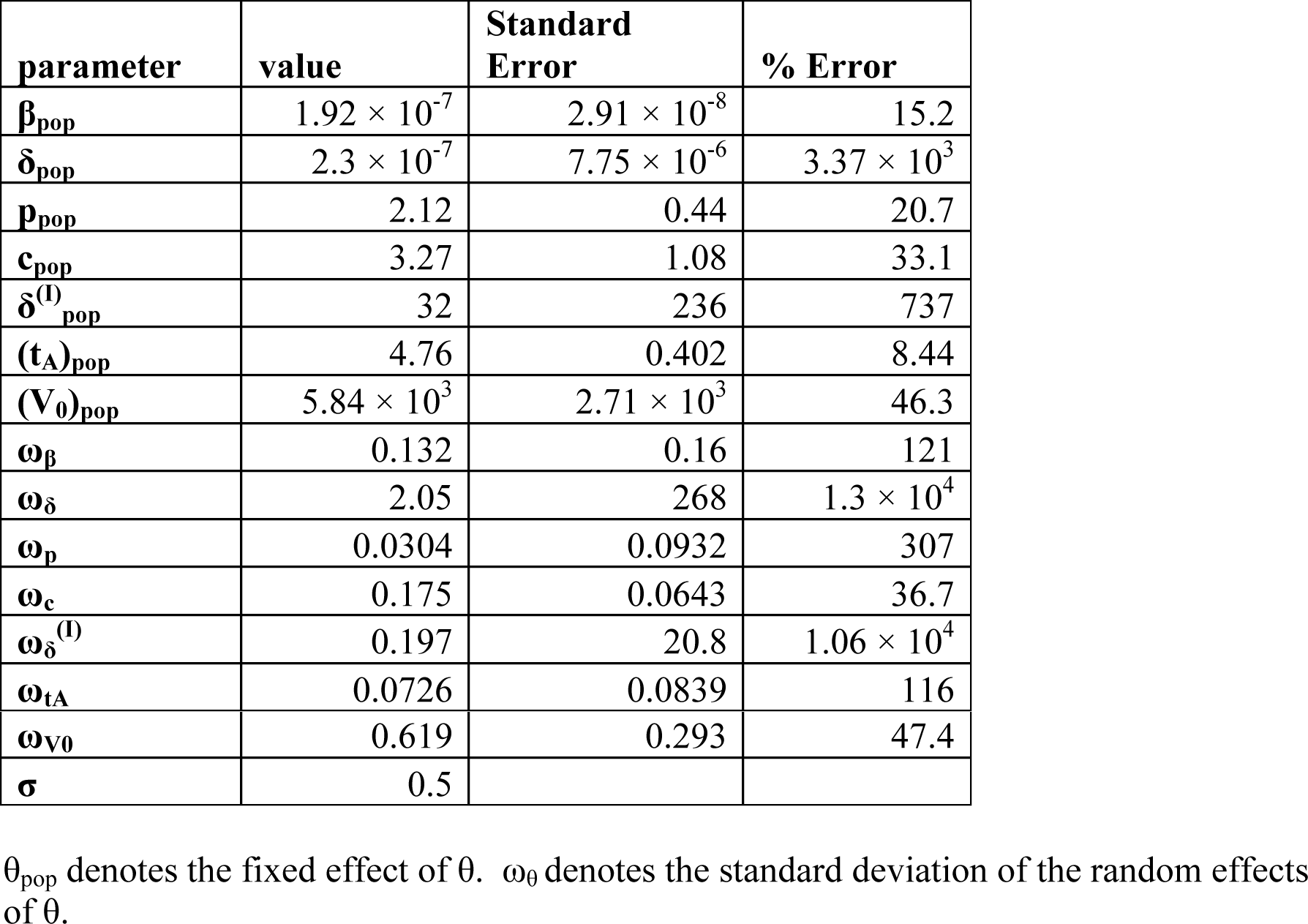
Parameter estimates for the data in Prince et al. where all (β, δ, p, c, δ^(I)^, t_A_, V_o_) the parameters were considered to be age dependent.

**Table S4:** Parameter estimates for the data in Prince et al. where β, c, δ^(I)^, and V_o_ were considered to be age dependent. The units of the parameters are the same as shown in Table II.

| parameter | value | Standard Error | % Error |
| --- | --- | --- | --- |
| $\beta_{\text{pop}}$ | $3.05 \times 10^{-7}$ | $2.80 \times 10^{-8}$ | 9.18 |
| $p_{\text{pop}}$ | 1.33 | 0.028 | 2.13 |
| $c_{\text{pop}}$ | 2.94 | 0.18 | 6.14 |
| $\delta^{(I)}_{\text{pop}}$ | 1153.9 | 1910.8 | 165.60 |
| $(t_A)_{\text{pop}}$ | 4.72 | 0.039 | 0.83 |
| $(V_0)_{\text{pop}}$ | 4504.5 | 1778.2 | 39.48 |
| $\omega_\beta$ | 0.00966 | 0.01101 | 113.96 |
| $\omega_c$ | 0.0049 | 0.0779 | 1580.58 |
| $\omega_{\delta^{(I)}}$ | 1.36 | 1.88 | 138.27 |
| $\omega_{V_0}$ | 0.693 | 0.263 | 37.90 |
| $\sigma$ | 0.5 | | |

**Table S5:** Parameter estimates for our dataset where β, c, δ^(I)^, and V_o_ were considered to be age dependent. The units of the parameters are the same as shown in Table II.

| parameter | value | Standard Error | % Error |
| --- | --- | --- | --- |
| $\beta_{\text{pop}}$ | $5.01 \times 10^{-7}$ | $8.68 \times 10^{-7}$ | 173.3 |
| $p_{\text{pop}}$ | 5.38 | 9.61 | 178.5 |
| $c_{\text{pop}}$ | 6.47 | 5.96 | 92.1 |
| $\delta^{(I)}_{\text{pop}}$ | 10.85 | 8.42 | 77.6 |
| $(t_A)_{\text{pop}}$ | 4.26 | 5.74 | 134.7 |
| $(V_0)_{\text{pop}}$ | 2.26 | 3.26 | 144.0 |
| $\omega_\beta$ | 0.065 | 0.141 | 216.6 |
| $\omega_c$ | 0.307 | 0.088 | 28.8 |
| $\omega_{\delta^{(I)}}$ | 0.038 | 0.267 | 699.3 |
| $\omega_{V_0}$ | 2.86 | 1.07 | 37.3 |
| $\sigma$ | 0.23 | | |

B. Age dependent estimates.
| **age (days)** | **$\beta$** | **$c$** | **$\delta^I$** | **$V_0$** |
| --- | --- | --- | --- | --- |
| **4** | $4.92 \times 10^{-7}$ | 4.74 | 10.7 | 0.161 |
| **10** | $4.99 \times 10^{-7}$ | 8.01 | 10.9 | 4.13 |
| **21+** | $5.00 \times 10^{-7}$ | 9.38 | 11.0 | 133 |
| **270+** | $5.2 \times 10^{-7}$ | 5.05 | 10.8 | 0.251 |

